# 3′UTR-mediated mRNA heterodimerization events pre-organize protein interactions to drive Wnt gene activation

**DOI:** 10.64898/2026.05.05.723075

**Authors:** Ting Cai, Nelly M. Cruz, Sudipto Basu, Richard M. White, Christine Mayr

## Abstract

Stem cell differentiation is controlled by transcription factors that are often encoded by mRNAs with highly conserved 3′UTRs. To identify 3′UTRs with the potential to drive differentiation, we partially deleted endogenous 3′UTRs. This altered stem cell differentiation efficiency in 7/10 cases without affecting encoded protein expression in 6/7 cases. For mechanistic investigation of abundance-independent regulatory roles of 3′UTRs, we focused on *CTNNB1*, an mRNA that encodes the essential Wnt co-activator β-catenin. *CTNNB1* 3′UTR deletion impairs zebrafish embryogenesis and Wnt gene induction during human stem cell differentiation. Long intermolecular 3′UTR-3′UTR interactions between Wnt transcription factor mRNAs and *CTNNB1* enable co-translational co-folding of their corresponding proteins. mRNA heterodimerization pre-allocates a portion of β-catenin molecules to the Wnt pathway and away from E-cadherin binding. This ensures protein co-localization with weaker partners, necessary for rapid and high Wnt gene induction. mRNA dimers exist before Wnt activation, indicating that they pre-organize future protein interactions.

## Introduction

Messenger RNAs (mRNAs) are essential for protein synthesis. The coding sequence (CDS) is translated into the protein sequence, whereas the roles of 3′ untranslated regions (3′UTRs) are less clear, as their functions seem to be diverse^1–7^. More than 2,700 human 3′UTRs have hundreds of nucleotides of exceptionally high sequence conservation^8,9^. In the highly conserved regions, nearly every nucleotide is conserved across vertebrate species (Fig. 1A), suggesting that base pairing and/or RNA structure are critical to mediate biological effects. To date, the functions of long, conserved 3′UTR regions are unknown.

**Figure 1.**
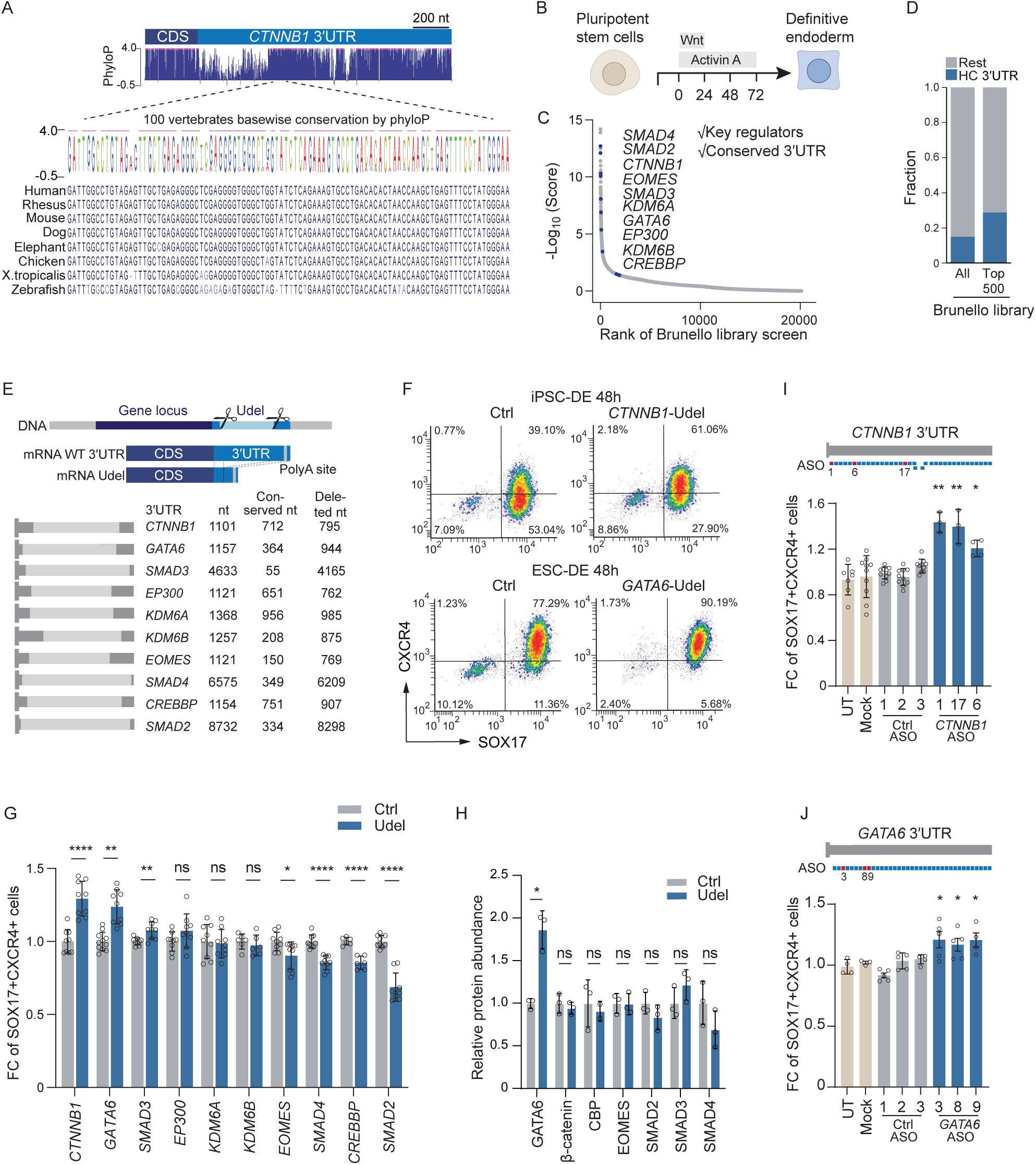
Endogenous 3′UTRs of transcriptional regulators modulate DE differentiation in a protein abundance-independent manner. **A.** UCSC genome browser tracks showing 100-way phyloP conservation of the *CTNNB1* 3′UTR. The zoomed-in region shows the nt sequences of selected vertebrate species. **B.** Schematic of the DE differentiation protocol. The Wnt pathway is induced through addition of the GSK3 inhibitor CHIR99021. **C.** Genes essential for ESCs to differentiate to DE. Shown is gene rank order from a genome-wide CRISPR screen. The 3′UTRs of the highlighted genes were deleted. **D.** HC 3′UTRs are significantly enriched among the top 500 essential genes (from C), compared with all genes in the Brunello library. X^2^ = 55.6; *P* < 0.0001. **E.** Schematic of 3′UTR loss-of-function approach. A genomic region that corresponds to partial deletion of a specific 3′UTR at the mRNA level is deleted from the human genome in pluripotent stem cells with CRISPR-Cas9 and a pair of guide RNAs. The locations of the guide RNAs ensure that the CDS and distal mRNA 3′ end processing elements are preserved. Shown is 3′UTR length, number of 3′UTR nt with phyloP ≥2 (considered conserved) and the number of nt deleted from endogenous 3′UTRs. **F.** Representative flow plots showing SOX17+ and CXCR4+ cells 48h post-DE induction in Ctrl and *CTNNB1* 3′UTR deletion (Udel) iPSCs (upper panel) or in Ctrl and *GATA6* Udel ESCs cells (bottom panel). **G.** Summary of SOX17+CXCR4+ double-positive cells obtained by flow cytometry analysis of ten different 3′UTR deletion candidates. For each candidate, three independent Ctrl clones were compared to three independent Udel clonal lines. Shown is mean ± SD from *N* = 3 independent experiments, each. For the *CREBBP* 3′UTR deletion, two independent clonal lines were generated. Welch’s *t*-test; *, *P* < 0.05; **, *P* < 0.01; ****, *P* < 0.0001; ns, not significant. **H.** Quantification of protein abundance measured by immunoblotting for the candidates with significant difference in DE differentiation efficiency from (G). Shown is mean ± SD from *N* = 3 clonal lines. Welch’s *t*-test, *P* = 0.016. Western blot data shown in Fig. S3A-G. **I.** Quantification of SOX17+CXCR4+ double-positive cells by flow cytometry 48h post-DE induction. iPSCs were untreated (UT), transfected with RNAiMAX only (mock), Ctrl ASOs, or ASO against the *CTNNB1* 3′UTR. Shown is mean ± SD from *N* = 3 independent experiments. Welch’s *t*-test was performed against the average of the Ctrl ASOs; *, *P* < 0.05; **, *P* < 0.01. **J.** As in (I), but HUES8 ESCs were transfected with Ctrl ASOs or ASO against the *GATA6* 3′UTR. Welch’s *t*-test; *, *P* < 0.05.

Massively parallel reporter assays using 3′UTR regions revealed a strong association between 3′UTR nucleotide content and repressive effects on mRNA and protein abundance^10,11^. Whereas GC-rich sequences were repressive, AU-rich 3′UTRs lacked such an effect. Moreover, a strong relationship between 3′UTR nucleotide content and sequence conservation was observed^10,11^. GC-rich 3′UTRs evolve rapidly and their mRNAs encode proteins with immune or metabolic roles, whereas AU-rich 3′UTRs are evolutionarily conserved and their mRNAs predominantly encode transcription factors and chromatin regulators^9,10^.

It was recently observed that highly conserved (HC) 3′UTRs are strongly enriched for proteins with long intrinsically disordered regions (IDRs)^9^. IDRs are protein segments with a lower fraction of hydrophobic amino acids compared with folded domains. Therefore, IDRs do not adopt a single stable conformation in isolation but sample a wide array of distinct conformational states^12,13^. For several transcriptional regulators with long IDRs, it was observed that full protein activity in cells requires the presence of HC 3′UTRs in mRNA templates during protein synthesis^9^. The HC 3′UTRs interact with the nascent chains and act as molecular chaperones to prevent co-translational misfolding of proteins with long IDRs^9,14,15^.

Gene functions are often studied through disruption of the endogenous gene locus^16^. However, similar loss-of-function experiments that only disrupt 3′UTRs are rare. So far, less than six endogenous 3′UTRs have been deleted in human cells or in mice^17–21^. The observed phenotypes range from a decreased kidney size to impaired thermogenesis and reduced stimulation-dependent neuronal activity. Here, we used CRISPR-Cas9 and pairs of guide RNAs to delete ten conserved endogenous 3′UTRs in factors expressed in human pluripotent stem cells and assessed the impact on stem cell differentiation efficiency. The results reveal that most tested 3′UTRs modulate stem cell differentiation efficiency in a protein abundance-independent manner.

We observed the strongest phenotype upon partial deletion of the *CTNNB1* 3′UTR. The *CTNNB1* mRNA encodes the β-catenin protein, which is best known as essential co-activator of the Wnt transcriptional program^22–24^. Wnt signaling is indispensable for embryonic development and cell fate decisions^25–27^. As Wnt transcription factors contain DNA-binding domains but lack activation domains, the Wnt transcriptional program is only induced when both β-catenin and the Wnt transcription factors bind to target gene promoters and enhancers, which then recruit co-activators, including p300/CBP and BCL9/BCL9L^23,24,28–30^. β-catenin also interacts with E-cadherin, which is essential for cell-cell adhesion. As E-cadherin and the Wnt transcription factors bind to the same site on β-catenin, their interaction is mutually exclusive^31^. However, it is currently unknown how the different pools of β-catenin, regulating adhesion or transcription are determined^23,32^.

We found here that the *CTNNB1* 3′UTR is required for full induction of the Wnt transcriptional program during human induced pluripotent stem cell (iPSC) differentiation and zebrafish embryogenesis. We discovered that the Wnt transcription factor mRNAs and *CTNNB1* form long intermolecular 3′UTR-3′UTR interactions that are necessary to drive the Wnt gene expression program. The mRNA interactions occur in 3′UTR regions with weak interaction to their own 3′UTRs, thus generating a strong propensity for intermolecular 3′UTR-3′UTR interactions. Heterodimerization of *CTNNB1* and the Wnt transcription factor mRNAs promotes co-translational protein co-folding between β-catenin and Wnt transcription factors. Our mechanistic investigation reveals that 3′UTR-driven protein co-folding provides quantitative and qualitative advantages over post-translational protein interactions, thus allowing rapid and strong Wnt target gene induction. Here, we reveal essential roles of HC 3′UTRs in coordinating signaling pathway-induced transcriptional programs, as sequential mRNA heterodimerization events pre-organize future protein interactions.

## Results

### HC 3′UTRs are enriched among genes essential for definitive endoderm differentiation

To study the functions of HC 3′UTRs (Fig. 1A), we set out to perform loss-of-function experiments. As mRNAs with HC 3′UTRs are enriched in transcription factors^9^, we used human pluripotent stem cell differentiation as experimental system. The combined activation of the Wnt and Activin A pathways differentiates pluripotent stem cells to definitive endoderm (DE) cells, which are the precursors of hepatocytes and pancreatic progenitor cells (Fig. 1B)^33^. Genes that are essential for DE differentiation were previously identified through a genome-wide CRISPR screen (Fig. 1C)^34^. Among the top 500 essential DE genes, mRNAs with HC 3′UTRs were two-fold enriched compared to their expected frequency among all genes in the CRISPR library (Fig. 1D)^9,34^. This observation suggests that HC 3′UTRs could play regulatory roles during DE differentiation.

To perform 3′UTR loss-of-function experiments, we used CRISPR-Cas9 and pairs of guide RNAs to delete genomic sequences, such that the resulting mRNA transcripts lack 68-95% of their 3’UTR sequence (Fig. 1E). The boundaries of the 3′UTR deletions (Udel) were chosen such that the CDS and distal mRNA 3′ end processing elements remained intact. The candidates for 3′UTR deletion were transcriptional regulators in the top 10% of genes essential for DE differentiation (Fig. 1C)^34^. These genes encode master transcription factors for DE differentiation, such as GATA6 and EOMES, co-activators, such as β-catenin, p300, CBP, the histone demethylases UTX and JMJD3, and signaling-induced transcription regulators, such as SMAD2, SMAD3, and SMAD4 (Table S1).

### 3′UTR deletions change stem cell differentiation efficiency in a protein abundance-independent manner

For each 3′UTR deletion, we generated at least three homozygous clones in human pluripotent stem cells, which were validated at the DNA and mRNA level (Fig. S1, Table S2). In cells with homozygous 3′UTR deletions, mRNA transcript expression was detected within their CDS but not within their respective 3′UTRs (Fig. S1C, S1D). We used DE differentiation efficiency as initial biological readout for potential 3′UTR effects, as the molecular consequences of 3′UTR deletions were unknown. We determined DE differentiation efficiency as previously described through the fraction of SOX17+CXCR4+ double-positive cells, measured by flow cytometry (Fig. 1F, S2)^33–35^. Overall DE differentiation efficiency in control (Ctrl) iPSCs was lower than in Ctrl HUES8 embryonic stem cells (ESC; Fig. 1F, S2B, S2C).

Comparing DE differentiation efficiency between Ctrl and 3′UTR deletion clones revealed a significant difference in SOX17+CXCR4+ cells in seven out of ten investigated genes, shown as fold change (FC) between Ctrl and Udel clones (Fig. 1F, 1G, S2D, S2E). We detected up to 30% more double-positive cells upon 3′UTR deletion of *CTNNB1*, *GATA6*, and *SMAD3* and observed significantly lower differentiation efficiency upon 3′UTR deletion of *EOMES*, *CREBBP*, *SMAD4*, and *SMAD2.* These findings suggest widespread 3′UTR regulatory roles during pluripotent stem cell differentiation.

Importantly, when comparing protein expression levels in Ctrl and 3′UTR deletion clones, measured by western blot or flow cytometry, we did not observe significant differences in 6/7 cases (Fig. 1H, S3A-I). This result indicates that most endogenous 3′UTR deletions did not significantly affect abundance levels of the mRNA-encoded proteins; rather, it suggests that most conserved 3′UTRs use regulatory mechanisms other than protein abundance to modulate stem cell differentiation. These observations are consistent with previously documented protein abundance-independent regulatory effects of AU-rich or HC 3′UTRs^2,9,10,18,36^.

### 3′UTR-targeting antisense oligonucleotides recapitulate the effects of the 3′UTR deletions

As we deleted regions in the DNA to generate 3′UTR mutant clones, the observed differences in DE differentiation efficiency could be caused through disruption of DNA rather than 3′UTR regulatory elements. To address this point, we repeated the DE differentiation in wildtype (WT) cells and blocked individual, 30-nt-long 3′UTR regions with antisense oligonucleotides (ASOs). The type of ASO used did not cause mRNA degradation but blocked sequence access^36,37^.

We tiled ASOs along the entire *CTNNB1* and *GATA6* 3′UTRs (Fig. S3J, Table S3). Remarkably, for each 3′UTR, we observed several ASOs that increased DE differentiation efficiency in a manner that was comparable to the corresponding 3′UTR deletion (Fig. 1I, 1J, S3K-N). These experiments confirmed that the observed changes in DE differentiation efficiency are caused by the mRNA 3′UTRs. Intriguingly, these results also revealed that the blocking of short 3′UTR regions can be sufficient to increase stem cell differentiation efficiency without manipulating the underlying DNA sequence.

### Deletion of the *GATA6* 3′UTR increases GATA6 protein levels and DE differentiation

Next, we set out to understand by which mechanisms mRNA 3′UTRs alter DE differentiation efficiency. GATA6 is one of the master transcription factors of DE differentiation^38,39^. *GATA6* transcript levels start to be expressed 18 hours (h) post-DE induction and peak at 48h (Fig. S4A)^40^. In 3′UTR deletion cells, at 24h post-DE induction, we observed more GATA6-positive cells, and these cells expressed higher GATA6 protein levels (Fig. S4B-D). GATA6 protein levels were significantly higher throughout DE differentiation in *GATA6* 3′UTR deletion cells compared to Ctrl cells (Fig. S4E). The higher GATA6 levels may explain the increase of CXCR4+SOX17+ double-positive cells from 77% to 90% (Fig. 1F, 1G).

For all other 3′UTR deletion candidates that showed differences in DE differentiation efficiency, protein abundance was not affected, thus making the mechanism of regulation unclear. To elucidate additional mechanisms by which HC 3′UTRs affect cellular processes, we focused on the *CTNNB1* 3′UTR, as its deletion had the strongest effect in modulating DE differentiation efficiency (Fig. 1F, 1G).

### *CTNNB1* 3′UTR deletion impairs Wnt target gene expression in a β-catenin abundance-independent manner

In the nucleus, β-catenin acts as essential co-activator of the Wnt signaling-induced transcriptional program (Fig. 2A)^22–24^. DE differentiation is induced through combined activation of the Activin A and Wnt pathways (Fig. 1B). In our differentiation protocol, Wnt is activated through addition of CHIR99021, a GSK3 inhibitor, at the start of DE differentiation^33,34^. Two well established Wnt target genes, *LEF1* and *AXIN2,* were transiently induced in Ctrl cells, but their induction was strongly impaired in *CTNNB1* 3′UTR deletion cells (Fig. 2B, 2C).

**Figure 2.**
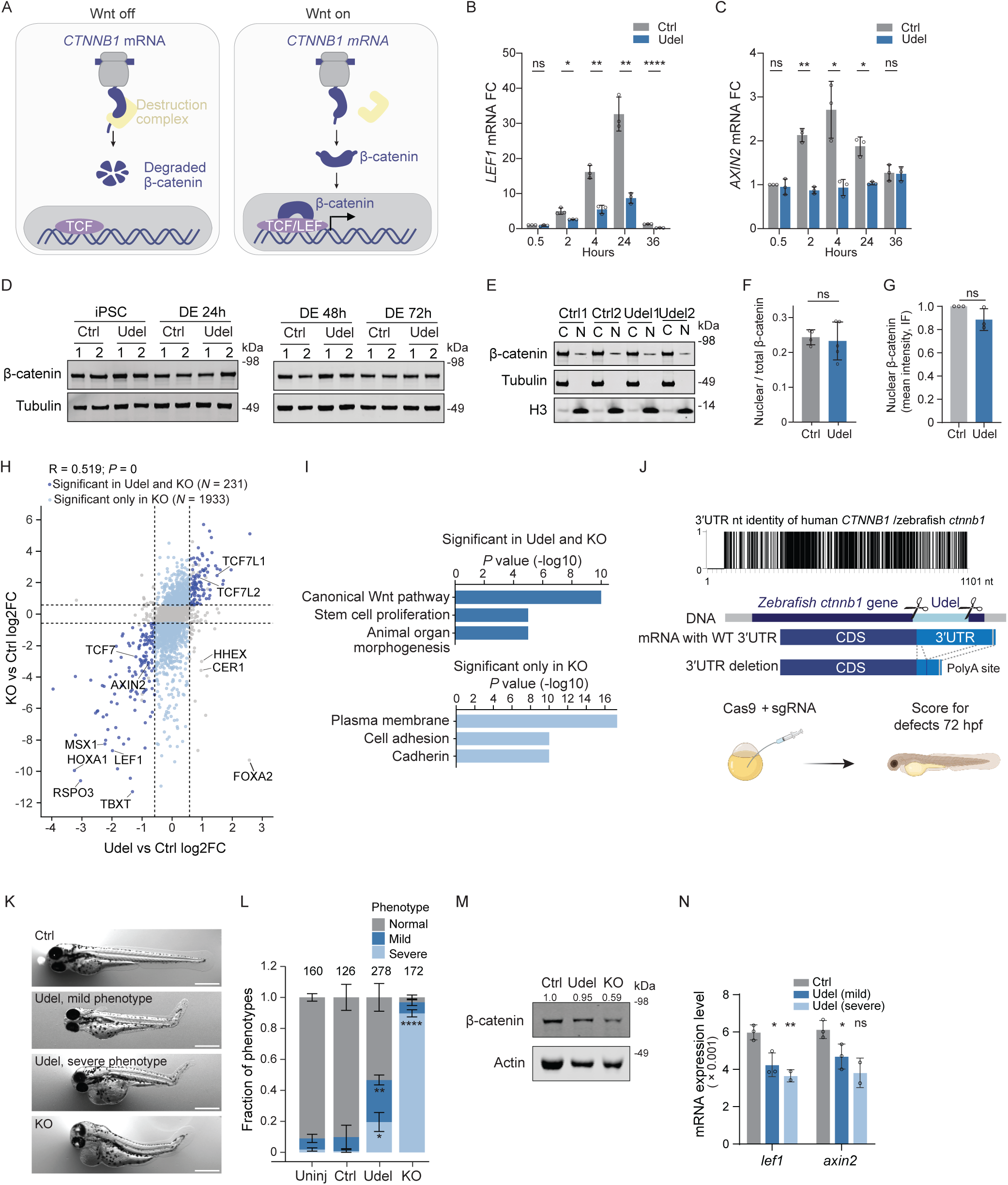
The *CTNNB1* 3′UTR is required for full induction of the Wnt gene expression program during iPSC differentiation and zebrafish embryonic development. **A.** Schematic of currently accepted model, by which β-catenin and the Wnt transcription factors activate the Wnt gene expression program. **B.** *LEF1* mRNA expression at the indicated timepoints, normalized to *GAPDH*. Shown is mean ± SD rom *N* = 3 independent experiments. Welch’s *t*-test; *, *P* < 0.05; **, *P* < 0.01; ****, *P* < 0.0001. **C.** As in (B), but *AXIN2* mRNA expression is shown. **D.** Immunoblot showing total β-catenin in Ctrl and Udel cells at the indicated timepoints. *N* = 2 clonal lines were examined. Tubulin serves as loading control. **E.** Immunoblot showing nuclear and cytoplasmic β-catenin at DE 24h loaded at 1:1 ratio. Tubulin serves as loading control for cytoplasmic fraction and H3 serves as loading control for nuclear fraction. **F.** Quantification of (E), shown as mean ± SD from *N* = 2 independent experiments where 2 or 3 clonal lines were examined. Welch’s *t-*test. **G.** Fluorescence intensity of nuclear β-catenin was quantified and shown as mean ± SD from 3 independent experiments. A representative image is shown in Fig. S4G. Welch’s *t*-test. **H.** Scatter plot showing log2FC in Udel versus Ctrl (x-axis) and log2FC in KO versus Ctrl (y-axis) at DE 24h of Wnt-responsive genes^27^. Genes with significant changes (log2|FC| > 0.58 and FDR < 0.05) in both Udel and KO are shown in dark blue (*N* = 231), whereas genes with significant change in KO only are shown in light blue (*N* = 1933). Dashed lines indicate a FC of 1.5. Selected genes are indicated. Pearson’s correlation coefficient is shown. **I.** Gene ontology analysis of genes colored in (H). Bonferroni-corrected *P* values are shown. **J.** Schematic of 3′UTR loss-of-function approach of the zebrafish *ctnnb1* gene. A genomic region of 776 bp was deleted using CRISPR-Cas9 and a pair of guide RNAs in fertilized eggs. At the mRNA level, this deletion results in partial deletion of the zebrafish *ctnnb1* 3′UTR. Embryonic defects are scored 72h after fertilization. Top panel, conserved nt between the human *CTNNB1* and the zebrafish *ctnnb1* 3′UTR. Each line denotes an identical nt. **K.** Representative images showing a normal zebrafish embryo, injected with a non-targeting guide RNA (Ctrl), mild and severe abnormalities observed after injection of a pair of guide RNAs that generate a *ctnnb1* 3′UTR deletion (Udel) and severe abnormalities after injection of a guide RNA targeting the *ctnnb1* coding sequence to generate a gene KO. Scale bar, 500 μm. **L.** Phenotype classification at 72h post-injection from experiment shown in (K). The total number of fish examined in each category is given. Shown is the mean fraction ± SD of the obtained phenotypes from three clutches obtained in two independent experiments. T-test for independent samples was performed; mild phenotype, uninjected (uninj) vs Udel, **, *P* =0.008; Uninj vs KO, ns; severe phenotype, uninj vs Udel, *, *P* = 0.046; uninj vs KO, ****, *P* = 4 x 10^−6^. **M.** Immunoblot showing total β-catenin obtained from zebrafish embryos at 72h post-injection. Four embryos were pooled for each sample. Actin was used as loading control. The numbers indicate relative protein abundance normalized to Actin in each sample. **N.** mRNA expression of *lef1* and *axin2* in zebrafish embryos 72 h post-injection, normalized to *eef1*. Shown is mean ± SD of *N* = 3 mRNA preparations that each contained four different embryos. Welch’s *t*-test; *, *P* < 0.05; **, *P* < 0.01.

According to the prevailing model (Fig. 2A), in the absence of Wnt signaling, the destruction complex co-translationally degrades β-catenin^41–43^. Wnt pathway activation allows β-catenin protein to be synthesized. It translocates to the nucleus, binds to the Wnt transcription factors and induces Wnt gene expression. Although Wnt pathway activation upregulates β-catenin protein levels in several models^44,45^, the increase in β-catenin protein level is small in ESCs^46^ and absent in the iPSCs we used. We did not observe increased β-catenin protein levels 24h after addition of CHIR99021 (Fig. 2D). Moreover, β-catenin protein abundance was not different between Ctrl and *CTNNB1* Udel cells at any of the four timepoints investigated during DE differentiation (Fig. 2D). We did not detect a difference in active β-catenin, which represents unphosphorylated β-catenin, between Ctrl and Udel cells (Fig. S4F). As Wnt activation is reported to promote translocation of β-catenin to the nucleus, we quantified β-catenin in the cytoplasmic and nuclear fractions 24h post-DE induction (DE 24h) and observed similar protein levels by western blot or immunofluorescence staining in Ctrl and Udel cells (Fig. 2E-G, S4G).

Taken together, our data indicate that *CTNNB1* 3′UTR deletion strongly reduced induction of Wnt target genes, such as *LEF1* and *AXIN2* during DE differentiation in a β-catenin protein abundance- and protein localization-independent manner. Our experimental system, which disrupts the *CTNNB1* 3′UTR but does not alter the endogenous β-catenin protein sequence or abundance, thus reveals a regulatory pathway of Wnt target gene induction that goes beyond simply regulating β-catenin protein levels.

### *CTNNB1* 3′UTR deletion impairs induction of canonical Wnt targets

To identify all the genes affected by the *CTNNB1* 3′UTR deletion, we performed RNA-seq at DE 24h. This highlighted two major groups of differentially expressed genes. Group (1) represented genes significantly affected by both knockout (KO) and Udel cells, whereas group (2) represented genes only affected by the *CTNNB1* KO (Fig. 2H, Table S4). By gene ontology analysis, group (1) represent canonical Wnt target genes, whereas the genes in group (2) are involved in cell adhesion and cadherin-mediated processes (Fig. 2I). Group (2) expression is strongly reduced in *CTNNB1* KO because β-catenin protein, which is essential for cell-cell adhesion in a Wnt pathway-independent manner, is absent.

For a few genes, we observed an expression change that was inconsistent between KO and Udel cells. Gene expression was reduced in KO cells, but was increased in Udel, compared with Ctrl cells. The genes include *CER1*, *HHEX*, and *FOXA2* (Fig. 2H) and represent crucial positive regulators of DE differentiation^47–49^. DE differentiation requires Activin A and low to intermediate Wnt signaling^50,51^. Activin A targets are upregulated and Wnt target gene expression is reduced but not absent in Udel cells. Taken together, the combination of both features provides an explanation for the, at first glance, surprising increase in DE differentiation efficiency observed in *CTNNB1* Udel cells (see Fig. 1F, 1G).

### *Ctnnb1* 3′UTR deletion causes embryonic abnormalities in zebrafish

To extend these results to the animal *in vivo*, we used zebrafish. The *CTNNB1* 3′UTR contains 1,101 nt and is highly conserved across vertebrates, including zebrafish (Fig. 1A). Zebrafish has two paralogs, called *ctnnb1* and *ctnnb2*. Intriguingly, the 3′UTR sequences of human and zebrafish *ctnnb1* are 61% identical (Fig. 2J, Table S1). Similar to human cells, we generated *ctnnb1* 3′UTR deletion and *ctnnb1* KOs by injecting Cas9 and guide RNAs into fertilized eggs (Fig. 2J). Disruption of canonical Wnt signaling in zebrafish embryos causes developmental phenotypes, because this pathway is central to axis formation and patterning. The reported phenotypes include shorter embryos and larvae, with abnormal body shape, and a characteristic bent tail^52^. *Ctnnb1* mutant zebrafish larvae generated with CRISPR-Cas9 exhibit disrupted tail development and abnormal body morphology. They were reported to have reduced head and eye size, and heart edemas^53^.

We assessed embryo morphology 72h after guide RNA injection. Embryos were classified into three groups of normal, mild, or severe abnormalities. If the embryos showed one of three changes, including shortened body length, bent tail or pericardial edema, their phenotypic abnormalities were considered mild, whereas if they had two or more changes, the abnormalities were considered severe. Whereas more than 90% of fish injected with Ctrl guide RNAs were phenotypically normal, 90% of embryos injected to generate *ctnnb1* KOs had severe abnormalities and phenocopied those previously reported (Fig. 2K, 2L)^53^. In the 3′UTR deletion group, 25% of embryos showed mild deformations, whereas 20% had severe abnormalities, despite mosaic editing (Fig. 2K, 2L, S4H-K).

As expected, β-catenin protein expression was reduced in *ctnnb1* KO embryos, but it was similar between *ctnnb1* Udel and Ctrl embryos (Fig. 2M). Despite similar β-catenin protein abundance, expression of the Wnt target genes *lef1* and *axin2* was significantly reduced in *ctnnb1* Udel compared with Ctrl embryos (Fig. 2N). These results suggest that 3′UTR-dependent regulation of Wnt target gene induction is conserved between human and zebrafish. Notably, deletion of a HC 3′UTR—even without significantly changing the level of expression of the mRNA-encoded protein—can cause severe embryonic abnormalities in a vertebrate animal.

### Wnt target gene expression requires β-catenin protein and the *CTNNB1* 3′UTR

Next, we investigated the mechanism by which the human *CTNNB1* 3′UTR controls Wnt target gene induction during iPSC differentiation. First, we performed a rescue experiment to ensure that the phenotype caused by the *CTNNB1* 3′UTR deletion is mediated by β-catenin protein (Fig. 3A). In *CTNNB1* KO cells, we knocked-in to the AAVS1 locus (KO-KI) a recombinant DNA containing the *CTNNB1* CDS and full-length 3′UTR, driven by a doxycycline-inducible promoter (Fig. S5A)^54,55^. Genes, whose expression was significantly reduced in *CTNNB1* Udel cells were largely rescued when a *CTNNB1* DNA including its 3′UTR was expressed in *CTNNB1* KO cells (Fig. 3B, 3C, S5B). The extent of rescue is consistent with a 22% reduction of β-catenin protein expression in KO-KI cells (Fig. 3D). This result indicates that both the *CTNNB1* 3′UTR and β-catenin protein need to be present for full Wnt target gene expression.

**Figure 3.**
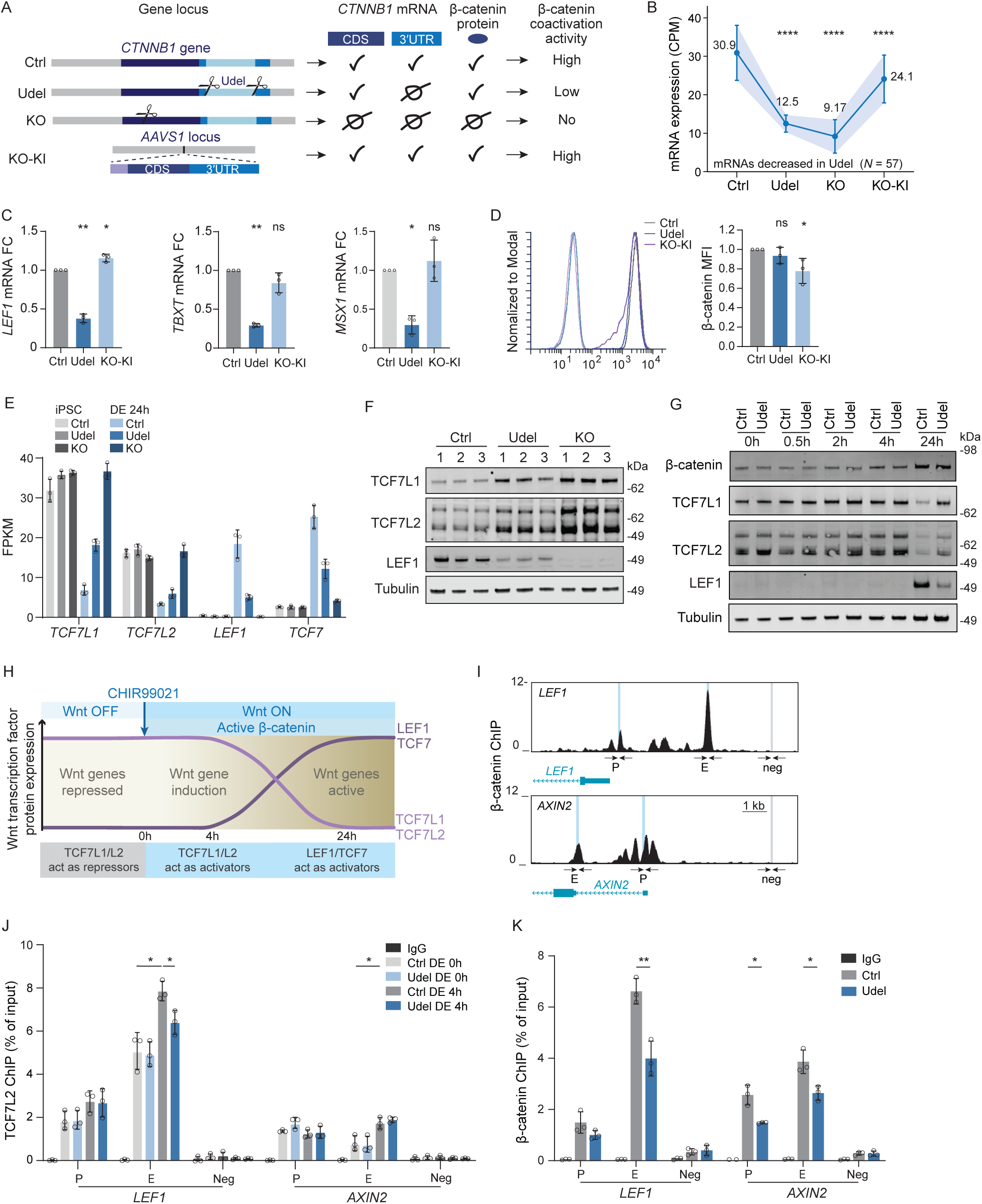
Upon Wnt signaling, the binding of β-catenin turns TCF7L1 and TCF7L2 into activating Wnt transcription factors. **A.** Schematic of rescue experiment. Shown is the human *CTNNB1* gene locus, the resulting mRNA transcript and protein product, together with β-catenin’s co-activation activity in the four investigated conditions. To obtain KO-KI cells, in *CTNNB1* KO cells, a recombinant DNA with the *CTNNB1* CDS and full-length 3′UTR was knocked-in to the AAVS1 locus. **B.** RNA-seq performed at DE 4h in the samples from (A). Gene expression (shown in counts per million, CPM) for all significantly reduced genes in *CTNNB1* Udel cells. Two-way Anova. ****, *P* < 0.0001. **C.** As in (B), but shown is mRNA FC for *LEF1, TBXT,* or *MSX1* normalized to *GAPDH*. Shown is mean ± SD from 3 independent experiments. Welch’s *t*-test. ***, *P* = 0.00036; **, *P* < 0.003; *, *P* = 0.031. **D.** β-catenin protein abundance measured at DE 4h by flow cytometry in samples from (A). Left, a representative flow plot. Unstained samples are shown in lighter colors. Right, β-catenin mean fluorescence intensity (MFI) as mean ± SD from 3 independent experiments. T-test for independent samples; *, *P* = 0.042. **E.** Fragments per kilobase of transcript per million mapped reads (FPKM) of Wnt transcription factor mRNAs obtained by RNA-seq at DE 24h. Shown is mean ± SD from 3 independent clonal lines. **F.** Immunoblot showing the indicated proteins at 24h post-DE induction in 3 clonal lines for Ctrl, Udel, and KO cells. Tubulin serves as loading control. **G.** As in (F), but protein expression at the indicated timepoints is shown. **H.** Schematic showing the regulation of Wnt transcription factor protein abundance before and after Wnt pathway activation. **I.** UCSC genome browser view of β-catenin chromatin immunoprecipitation (ChIP) sequencing at the *LEF1* and *AXIN2* gene loci. P, promoter; E, enhancer; neg, negative ctrl regions. **J.** TCF7L2 ChIP qRT-PCR performed at the genomic loci from (I) before (0h) and 4h post-DE induction. Shown is mean ± SD from 3 independent experiments. Welch’s *t*-test; *, *P* < 0.05. **K.** As in (J), but β-catenin ChIP qRT-PCR is shown at DE 4h. Welch’s *t*-test; *, *P* < 0.05; **, *P* < 0.01.

### At early timepoints of Wnt activation, presence of β-catenin turns TCF7L1 and TCF7L2 into activating Wnt transcription factors

Wnt target gene induction requires binding of both β-catenin and Wnt transcription factors to promoters and enhancers of Wnt-responsive genes^27,56^. The Wnt transcription factors TCF7L1 and TCF7L2 are expressed in iPSCs and their mRNA and protein levels strongly decrease within the first 24h post-DE induction (Fig. 3E-G). The Wnt transcription factors LEF1 and TCF7 show the opposite expression profile: they are not present in iPSCs but become the dominant factors at 24h (Fig. 3E-G). According to the prevailing model, the activating transcription factors LEF1/TCF7 replace the inhibitory Wnt transcription factors TCF7L1/TCF7L2 during the switch from a Wnt-off to a Wnt-on state^57^. This replacement does not occur in *CTNNB1* KO cells and happens only partially in *CTNNB1* 3′UTR deletion cells (Fig. 3E, 3F). This result shows that the *CTNNB1* 3′UTR is required for full LEF1 induction and for full TCF7L1/TCF7L2 protein repression during the first 24h of DE differentiation.

We turned to earlier timepoints to study the most direct effects of the *CTNNB1* 3′UTR. Within the first 4h of DE induction, protein expression levels of β-catenin, TCF7L1 and TCF7L2 were unaltered across the timepoints and were not different between Ctrl and Udel cells (Fig. 3G). Nevertheless, induction of *LEF1* and *AXIN2* mRNAs was strongly reduced (Fig. 2B, 2C). *LEF1* is a highly sensitive Wnt target gene and encodes one of the activating Wnt transcription factors during stem cell differentiation^27^. Wnt-induced LEF1 expression generates a positive feed-forward loop, which is necessary for sustained expression of Wnt target genes^22–24,58^. As LEF1 protein is not detectable within the first 4h of DE differentiation (Fig. 3G), but Wnt targets are transcribed, our data indicate that binding of β-catenin turns TCF7L1 and TCF7L2 into activating transcription factors at early timepoints after Wnt activation (Fig. 3H)^59^. In our experimental system, expression levels of all participating proteins were unaltered within the first 4h post-DE induction. This finding revealed a protein abundance-independent regulatory switch during early stages of Wnt activation that is impacted by the *CTNNB1* 3′UTR. All subsequent experiments were performed during this time frame.

### *CTNNB1* 3′UTR deletion reduces β-catenin occupancy at Wnt gene enhancers

Wnt gene induction requires binding of Wnt transcription factors at their promoters and enhancers. Chromatin immunoprecipitation (ChIP) of TCF7L2 at the *LEF1* and *AXIN2* gene loci showed increased TCF7L2 chromatin occupancy at DE 4h in Ctrl cells^27^, consistent with an activating role of TCF7L2 at early stages after Wnt induction (Fig. 3H-J). However, β-catenin ChIP showed significantly lower chromatin occupancy at the *LEF1* and *AXIN2* gene loci in Udel versus Ctrl cells (Fig. 3K). Reduced β-catenin co-activator levels at Wnt gene enhancers are consistent with reduced induction of *LEF1* and *AXIN2*, but the mechanism by which the *CTNNB1* 3′UTR increases β-catenin’s chromatin occupancy without changing overall protein levels was unclear.

### The *CTNNB1* 3′UTR increases co-localization between *CTNNB1* and Wnt transcription factor mRNAs

According to the currently accepted model, Wnt transcription factors and β-catenin translocate to the nucleus independently and subsequently bind to each other post-translationally at promoters and enhancers (Fig. 4A)^22–24^. In addition to post-translational binding, proteins can also interact co-translationally, which is estimated to occur in 40% of human proteins^60,61^. Co-translational interaction enables proteins to form stable heterodimers earlier, during protein synthesis. In human cells, co-translational assembly has mostly been studied in HEK293T cells, where co-translational heterodimerization of β-catenin and LEF1 was detected^61^.

**Figure 4.**
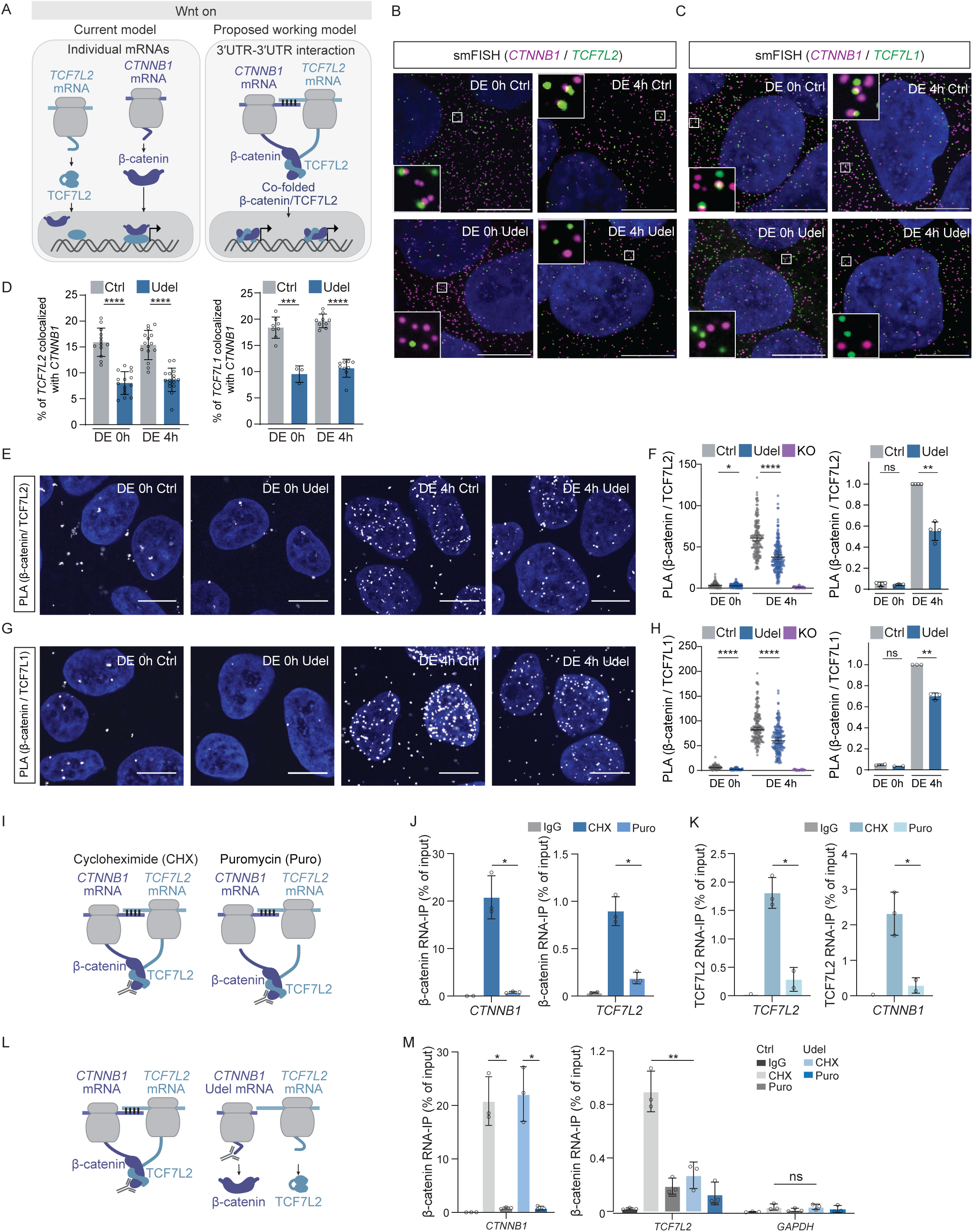
The *CTNNB1* 3′UTR is necessary for co-translational protein co-folding of β-catenin/TCF7L2 and β-catenin/TCF7L1. **A.** Current and proposed working model for Wnt signaling-induced target gene activation mediated by β-catenin and the Wnt transcription factor TCF7L2. **B.** Representative images of smRNA-FISH of endogenous *CTNNB1* and endogenous *TCF7L2* in iPSCs at 0h and 4h post-DE induction. The white square indicates the zoomed-in region. Scale bar, 10 μm. **C.** As in (B), but shown are *CTNNB1* and *TCF7L1* mRNAs. **D.** Quantification of experiments shown in (B) and (C). Left panel, percent of co-localized endogenous *TCF7L2* with *CTNNB1* is shown as mean ± SD from 2 independent experiments. Welch’s *t*-test. DE 0h, ****, *P* = 2 x 10^−8^; DE 4h, ****, *P* = 8 x 10^−8^. Right panel, percent of co-localized endogenous *TCF7L1* with *CTNNB1.* Welch’s *t*-test. DE 0h, ***, *P* = 0.0008; DE 4h, ****, *P* = 2x 10^−9^. At least 70 cells were analyzed in each experiment. See S5C and S5D for the number of foci. **E.** Representative images of protein PLA foci of endogenous β-catenin and endogenous TCF7L2 at 0h and 4h post-DE induction. DAPI was used for nuclear staining. Scale bar, 10 μm. **F.** Left panel shows mean ± SEM from 2 independent experiments, described in (E). 0h, Ctrl, *N* = 136, Udel, *N* = 146, Welch’s *t*-test, *, *P=*0.049; DE 4h, Ctrl, *N* = 153; Udel, *N* = 186; KO, *N* = 49, ****, *P* = 4 x 10^−21^. Right panel shows mean ± SD from 4 independent experiments. Welch’s *t*-test; **, *P* = 0.002. **G.** As in (E), but shown is PLA of endogenous β-catenin and endogenous TCF7L1 at 0h and 4h post-DE induction. **H.** Left panel shows mean ± SEM from 2 independent experiments, described in (E). 0h, Ctrl, *N* = 83; Udel, *N* = 66, *, *P =* 0.049; DE 4h, Ctrl, *N* = 172; Udel, *N* = 155; KO, *N* = 46, Welch’s *t*-test ****, *P* < 3 x 10^−13^. Right panel shows mean ± SD from 3 independent experiments. Welch’s *t*-test; **, *P* = 0.0037. **I.** Schematic depicting approach to test co-translational protein complex assembly between β-catenin and TCF7L2 nascent chains by RNA immunoprecipitation assay (RNA-IP). **J.** β-catenin RNA-IP, followed by qRT-PCR for *CTNNB1* and *TCF7L2* mRNAs at 2h post-DE induction in Ctrl cells. Shown is mean ± SD from 3 independent experiments. Welch’s *t*-test; *, *P* < 0.05. **K.** As in (J), but TCF7L2 RNA-IP followed by qRT-PCR is shown. **L.** Schematic of co-translational protein complex assembly between β-catenin and TCF7L2 nascent chains in Ctrl and Udel cells. **M.** As in (J), but shown for Ctrl and Udel cells. Welch’s *t*-test; **, *P* < 0.01.

To explain our results, we hypothesized the following working model: A 3′UTR-3′UTR interaction between the *CTNNB1* and *TCF7L2* mRNAs brings the ribosomes that translate β-catenin and TCF7L2 into proximity to enable their co-translational protein heterodimerization. The co-folded β-catenin/TCF7L2 dimer does not dissociate and is ready to induce Wnt target genes (Fig. 4A). *CTNNB1* 3′UTR deletion disrupts co-translational protein co-folding and represents a scenario where only individual protein molecules are synthesized, which interact post-translationally, a scenario that is less effective with respect to Wnt target gene induction (Fig. 4A).

To test our working model experimentally, we examined whether mRNA co-localization, protein co-localization, and co-translational protein interaction require the *CTNNB1* 3′UTR. We performed single molecule (sm) RNA-FISH on the endogenous mRNAs before and 4h post-DE induction. In all conditions, *CTNNB1* mRNA is present at much higher abundance than the Wnt transcription factor mRNAs *TCF7L1* and *TCF7L2*, suggesting that they are the limiting components (Fig. S5C, S5D). We observed substantial co-localization between *CTNNB1* and *TCF7L2* or between *CTNNB1* and *TCF7L1* mRNAs and in both cases, the interaction was reduced to approximately 50% in *CTNNB1* Udel cells. In Ctrl cells, 15% of *TCF7L2* and 18% of *TCF7L1* mRNAs colocalized with *CTNNB1* (Fig. 4B-D). Surprisingly, the co-localization between *CTNNB1* and the Wnt transcription factor mRNAs already existed before Wnt pathway induction, and its magnitude was unaffected by Wnt signaling (Fig. 4B-D).

### The *CTNNB1* 3′UTR increases protein co-localization of β-catenin and TCF7L1 or TCF7L2

Next, we used a proximity ligation assay (PLA) to examine co-localization of the endogenous proteins. Protein proximity within 40 nm is indicated by puncta formation^62^. Before Wnt induction, β-catenin/Wnt transcription factor complexes were not present (Fig. 4E-H), consistent with β-catenin degradation by the destruction complex in the absence of Wnt signaling^42^.

At DE 4h, however, we detected robust protein co-localization between β-catenin and TCF7L2, which was reduced to 50% in Udel versus Ctrl cells (Fig. 4E, 4F). For β-catenin/TCF7L1 protein co-localization, we observed a 30% reduction in PLA puncta in Udel cells (Fig. 4G, 4H). These results demonstrate that the *CTNNB1* 3′UTR promotes nuclear protein co-localization between β-catenin and the two Wnt transcription factors at DE 4h.

This model predicts co-translational heterodimerization, indicating that the two proteins should already interact in the cytoplasm. We performed β-catenin co-immunoprecipitation (co-IP) from the cytoplasmic fraction and detected robust interaction with either TCF7L1 or TCF7L2 in Ctrl cells 4h post-DE induction, but a strongly reduced interaction in Udel cells (Fig. S5E), which is consistent with the proposed model.

### The *CTNNB1* 3′UTR is necessary for co-translational co-folding of β-catenin/TCF7L1 and β-catenin/TCF7L2

To obtain direct evidence of co-translational protein heterodimerization in Ctrl cells, we performed a previously described RNA immunoprecipitation (RNA-IP) experiment at 4h post-DE induction^63,64^. In case of co-translational protein heterodimerization, β-catenin RNA-IP should pull down its own mRNA as well as *TCF7L2* mRNA, as they are connected to the β-catenin nascent chain through their respective ribosomes (Fig. 4I). To rule out direct β-catenin binding to these mRNAs, the RNA-IP was repeated in the presence of puromycin, which dissociates the nascent chains from the ribosomes (Fig. 4I)^63,64^. At 2h post-DE induction, β-catenin RNA-IP pulls down *CTNNB1* and *TCF7L2* mRNAs. Their pull down is strongly reduced upon puromycin treatment, indicating that β-catenin and TCF7L2 nascent chains interact (Fig. 4J). A similar result was obtained when an antibody against TCF7L2 was used for RNA-IP (Fig. 4K), demonstrating co-translational heterodimerization of β-catenin and TCF7L2 nascent chains.

To examine whether co-translational heterodimerization is 3′UTR-dependent, we repeated the experiment in *CTNNB1* Udel cells; this showed strongly reduced pull down of *TCF7L2* mRNA without reducing pull down of *CTNNB1* mRNA (Fig. 4L, 4M). This result demonstrates that co-translational protein co-folding of β-catenin and TCF7L2 requires the *CTNNB1* 3′UTR. Similar results were obtained for 3′UTR-dependent co-folding between β-catenin and TCF7L1 (Fig.

S5F).

Before Wnt induction, we detected mRNA co-localization without protein co-localization by smRNA-FISH and PLA (Fig. 4D, 4F, 4H). When performing β-catenin RNA-IP before Wnt induction, *TCF7L2* mRNA was not pulled down (Fig. S5G, S5H). This experiment serves as further control showing that β-catenin RNA-IP captures mRNAs associated through co-translational protein assembly, but not pre-associated mRNAs that have not formed a detectable nascent chain complex before Wnt activation.

To directly observe nascent chain interaction of β-catenin and TCF7L2 in live cells, we used the SunTag and MoonTag system (Fig. S5I)^65,66^. The two tags consist of repeated peptides recognized by respective co-expressed antibodies or nanobodies, thus allowing single-molecule live cell imaging of specific nascent chains. The SunTag was fused with *CTNNB1* CDS and 3′UTR and the MoonTag was fused to *TCF7L2* CDS and 3′UTR and expressed in HEK293T cells. With this system, we observed cytoplasmic co-localization of β-catenin and TCF7L2 at 2h after Wnt induction in live cells, suggesting co-translation (Fig. S5J).

### Deletion of the *TCF7L2* 3′UTR reduces β-catenin/TCF7L2 protein co-localization

To obtain additional evidence for the proposed intermolecular 3′UTR-3′UTR interaction between the *CTNNB1* and *TCF7L2* mRNAs, we partially deleted the 3′UTR of *TCF7L2*. The genomic deletion removed 1,430 nt of the *TCF7L2* 3′UTR without disrupting the CDS or distal mRNA 3′ end processing elements (Fig. 5A, S6A, S6B). The *TCF7L2* 3′UTR deletion did not affect TCF7L2 protein levels in iPSCs or at DE 4h (Fig. 5B, S6C). PLA performed in *TCF7L2* Udel cells showed that β-catenin/TCF7L2 protein PLA puncta were reduced by 40%, which was similar to the reduction observed in *CTNNB1* Udel cells (Fig. 5C). This result strongly supports that a 3′UTR-3′UTR interaction between the *CTNNB1* and *TCF7L2* mRNAs is required for the formation of β-catenin/TCF7L2 co-folded protein dimers.

**Figure 5.**
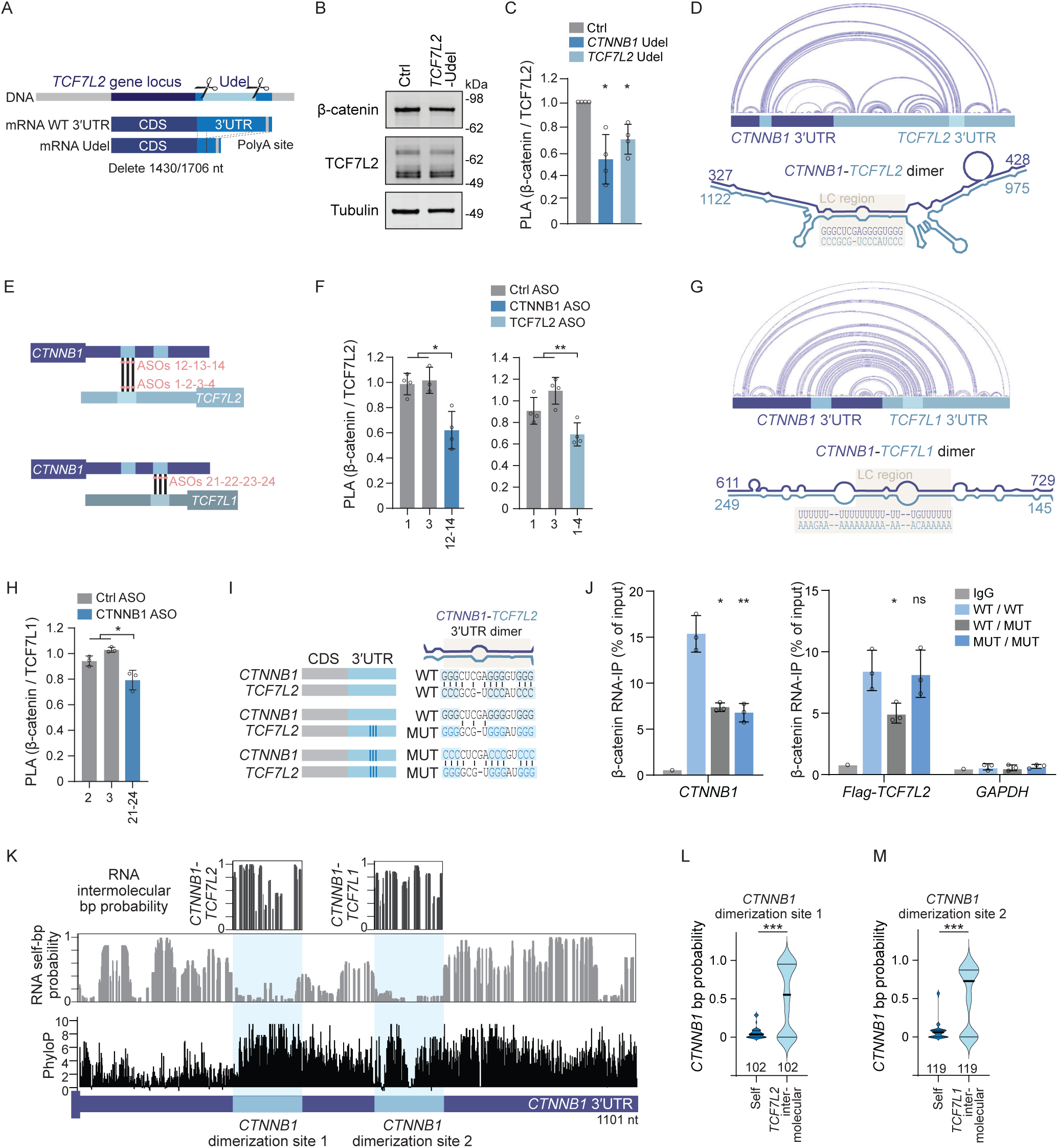
Accessible regions in the *CTNNB1* 3′UTR form intermolecular 3′UTR-3′UTR interactions with Wnt transcription factor mRNAs. **A.** Schematic of *TCF7L2* 3′UTR deletion. Shown as in Fig. 1E. **B.** Immunoblots showing TCF7L2 and β-catenin in Ctrl and *TCF7L2* Udel cells at DE 4h. Tubulin was used as loading control. **C.** Quantification of β-catenin/TCF7L2 PLA signal at DE 4h as mean ± SD from 4 independent experiments. Welch’s *t*-test; *, *P* < 0.02. **D.** RNA bifold prediction of mRNA heterodimerization between the *CTNNB1* and *TCF7L2* 3′UTRs. The major dimerization sites are highlighted, and the predicted structure of the 3′UTR dimer is shown below. The boundary nt of the 3′UTR dimer are indicated and the nt of participating low complexity (LC) regions are shown. **E.** Schematic of the intermolecular 3′UTR-3′UTR interactions between *CTNNB1* (dark blue) and *TCF7L2* (light blue; top panel). The mRNA heterodimerization sites are indicated with lighter colors. The vertical lines indicate RNA-RNA interactions, which are blocked by the indicated ASOs. The taller blocks in the mRNAs indicate the CDS, which is located upstream of the 3′UTR. Bottom panel shows 3′UTR-3′UTR interaction between *CTNNB1* and *TCF7L1*. **F.** Quantification of β-catenin/TCF7L2 PLA signal at DE 4h after transfection of ASOs in WT iPSCs. Shown is mean ± SD from at least 4 independent experiments. Welch’s *t*-test; **, *P* < 0.009. **G.** As in (D), but RNA bifold prediction of mRNA heterodimerization between the *CTNNB1* and *TCF7L1* 3′UTRs is shown. **H.** As in (F), but shown for the β-catenin/TCF7L1 PLA signal. Shown is mean ± SD from 3 independent experiments. Welch’s *t*-test; *, *P* = 0.029. **I.** Schematic for disruption of the *CTNNB1*-*TCF7L2* 3′UTR dimerization site using the indicated point mutations within the LC region. Intermolecular base pairing is rescued through introduction of compensatory mutations in the interacting 3′UTR. **J.** β-catenin RNA-IP, performed in HEK293T cells containing the constructs from (I). qRT-PCR for stably expressed *CTNNB1* mRNA and transfected *Flag-TCF7L2* mRNA at 6h post CHIR99021 treatment. Shown is mean ± SD from 3 independent experiments. Welch’s *t*-test; *, *P* < 0.048. **, *P =* 0.007. *GAPDH* mRNA is shown as control. **K.** Shown are the two *CTNNB1* 3′UTR dimerization sites (highlighted in light blue) together with phyloP score distribution and RNA self-base pairing (bp) probability for each nucleotide in the *CTNNB1* 3′UTR. Top panel, the intermolecular RNA bp probabilities of *CTNNB1* and the indicated Wnt transcription factor mRNAs within the dimerization sites is shown. **L.** Base pairing (bp) probability within the *CTNNB1* dimerization site 1 is shown for *CTNNB1* RNA self-interaction and the *CTNNB1-TCF7L2* intermolecular interaction. Wilcoxon test, *P* = 1.5 x 10^−11^. **M.** As in (L), but RNA bp probability within the *CTNNB1* dimerization site 2 is shown for *CTNNB1* self-interaction and the *CTNNB1-TCF7L1* intermolecular interaction. Wilcoxon test, *P* = 2.8 x 10^−13^.

### Blocking the formation of 3′UTR dimers through ASOs reduces protein co-localization

Next, we set out to map the 3′UTR-3′UTR interaction sites. The RNA bimolecular folding (bifold) program co-folds two given RNAs and predicts their interacting nucleotides (nt)^67^. For the 3′UTRs of *CTNNB1* and *TCF7L2,* RNA bifold predicts a 102-nt-long RNA interaction, of which 64 base pair intermolecularly (Fig. 5D, S6D, Table S5). *CTNNB1* ASOs 12-14 are predicted to block this interaction (Fig. 5E). We performed PLA in WT cells, either transfected with Ctrl ASOs or ASOs 12-14, and observed a 50% reduction in β-catenin/TCF7L2 interaction in ASO 12-14-treated cells (Fig. 5F). A similar reduction was observed in WT cells transfected with ASOs against the corresponding interacting region in the *TCF7L2* 3′UTR (ASOs 1-4; Fig. 5E, 5F). Addition of the ASOs did not affect protein abundance of β-catenin or TCF7L2 (Fig. S6E, S6F). Remarkably, the 3′UTR-blocking ASOs fully mimicked the 3′UTR deletion effects on protein co-localization (Fig. 5C, 5F), showing that the *CTNNB1-TCF7L2* mRNA heterodimer is both necessary and sufficient for 3′UTR-dependent co-folding of β-catenin and TCF7L2 in WT cells.

RNA bifold predicts a similar RNA interaction between the *CTNNB1* and *TCF7L1* 3′UTRs. This interaction spans 119 nt and has 76 intermolecular base pairs (Fig. 5G, S6G). *CTNNB1* ASOs 21-24 are predicted to disrupt it (Fig. 5E). Adding these ASOs to WT cells reduced β-catenin/TCF7L1 protein PLA puncta by 30% compared to Ctrl ASOs, and mimicked the results obtained in *CTNNB1* 3′UTR deletion cells (Fig. 4H, 5H). Next, we investigated whether the *CTNNB1* 3′UTR contains two distinct interaction sites, bound by either *TCF7L1* or *TCF7L2*, or whether blocking one region is sufficient to disrupt both interactions (Fig. 5D, 5G). We observed that the *CTNNB1* 3′UTR contains two distinct binding sites (Fig. 5E). Blocking the *CTNNB1-TCF7L2* interaction with ASOs 12-14 did not block the *CTNNB1-TCF7L1* interaction (Fig. S6H); similarly, blocking the *CTNNB1-TCF7L1* interaction with ASOs 21-24 did not block the *CTNNB1-TCF7L2* interaction (Fig. S6I).

### Evidence for direct *CTNNB1-TCF7L2* 3′UTR dimerization

To obtain definitive evidence for a 3′UTR-3′UTR interaction between *CTNNB1* and *TCF7L2*, we examined whether 3′UTR point mutations disrupt co-translational protein co-folding and whether compensatory mutations in the interacting 3′UTR can rescue protein co-folding (Fig. 5I). We stably expressed *CTNNB1* CDS and 3′UTR in *CTNNB1* KO HEK293T cells and transiently transfected *TCF7L2* constructs containing its CDS and 3′UTR. We observed similar protein levels of β-catenin across conditions (Fig. S6J, S6K).

β-catenin RNA-IP revealed reduced co-translational protein co-folding when one side of the 3′UTR heterodimer was disrupted, which was restored with compensatory mutations on the other side of the 3′UTR heterodimer (Fig. 5J). This experiment demonstrates direct interaction between the *CTNNB1* and *TCF7L2* mRNAs, which is necessary for their corresponding nascent chain interactions and co-translational co-folding of their respective proteins.

Direct mRNA interaction is further supported by psoralen-based chemical crosslinking, performed in iPSCs. Psoralen is a photoactivatable RNA crosslinker and has previously been used to detect direct RNA-RNA interactions in cells^68,69^. After RNA-RNA crosslinking, we used biotinylated probes to pull down specific mRNAs (Fig. S6L). Biotinylated probes against *CTNNB1* enriched *CTNNB1* mRNA ∼500-fold compared with a control probe (Fig. S6M). In Ctrl and Udel cells, the enrichment of *CTNNB1* mRNA was similar, but enrichment of interacting *TCF7L2* mRNA was diminished in Udel versus Ctrl cells (Fig. S6N), suggesting that the endogenous *CTNNB1* and *TCF7L2* 3′UTRs directly interact in iPSCs.

### Weak RNA self-interaction promotes formation of intermolecular RNA interactions

To learn what RNA features promote formation of intermolecular 3′UTR-3′UTR interactions, we determined the minimum free energy structure predicted for the *CTNNB1* 3′UTR using RNAfold^70,71^. RNAfold color-codes each nt of a given RNA according to its predicted self-base pairing probability (Fig. S6O). After extracting the numerical values, we observed that the heterodimerization region in the *CTNNB1* 3′UTR has very weak self-interaction potential (Fig. 5K). In contrast, the RNA intermolecular interactions with *TCF7L1* or *TCF7L2* are predicted to be strong (Fig. 5K-M, S6P). According to thermodynamics, an RNA region that binds weakly to itself has a high propensity to be replaced by a new interaction, if more energy is released from forming the new interaction than what was spent during disruption of the existing intramolecular interactions^72^. Our data suggest that intrinsically unpaired RNA regions are hotspots for the formation of intermolecular 3′UTR-3′UTR interactions.

### The protein-protein interaction between β-catenin and E-cadherin is 3′UTR independent

As β-catenin binds to proteins other than the Wnt transcription factors, we set out to test whether additional β-catenin protein interactions require the *CTNNB1* 3′UTR. One of the major β-catenin binding partners is E-cadherin and this interaction is essential for cell-cell adhesion^32^. According to PLA, the interaction between β-catenin and E-cadherin is abundant, but the protein co-localization does not require the *CTNNB1* 3′UTR (Fig. 6A, 6B). Also, the two encoding mRNAs co-localize in a 3′UTR-independent manner (Fig. 6C, 6D, S7A).

**Figure 6.**
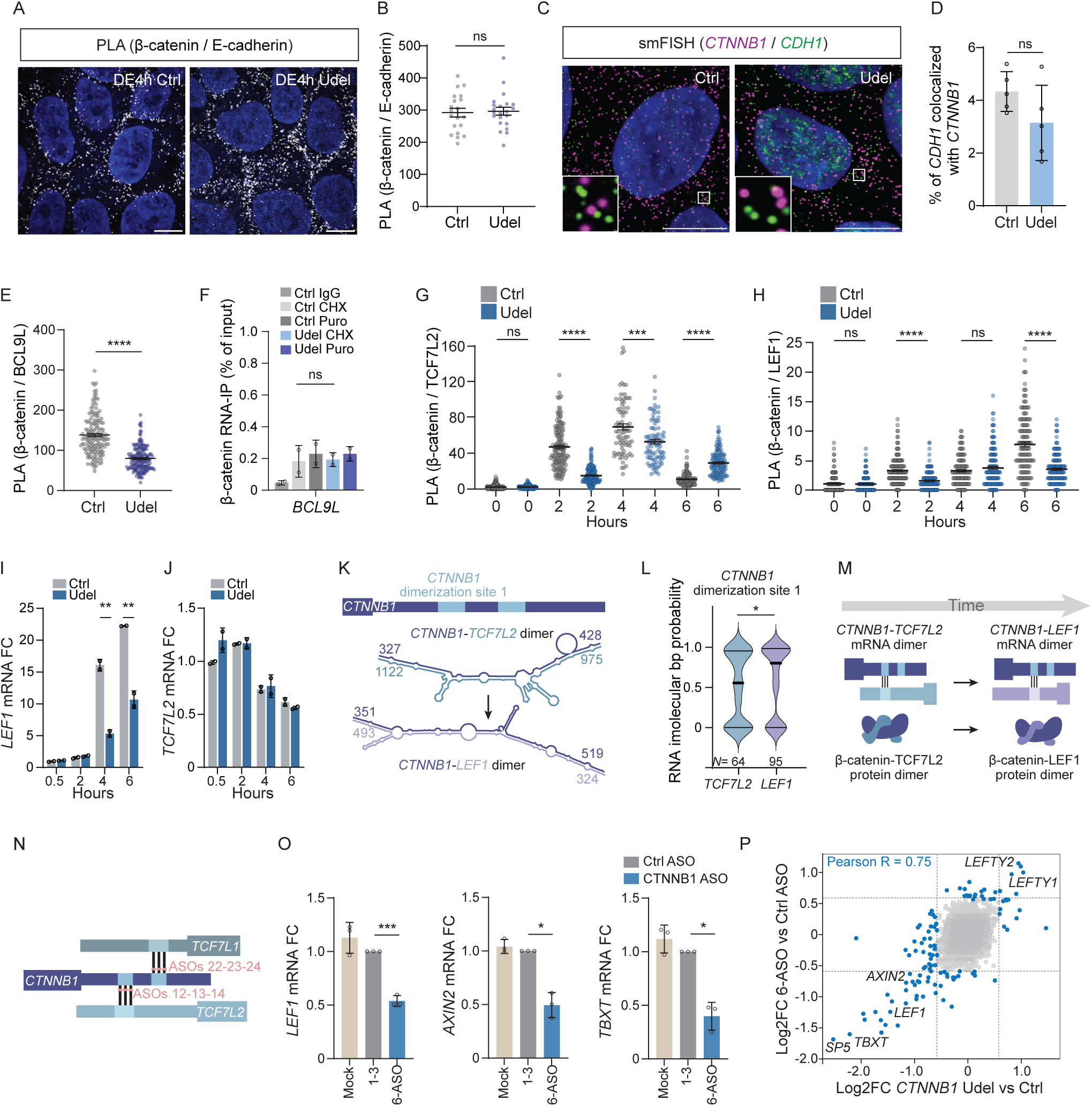
ASO-mediated blocking of *CTNNB1* mRNA heterodimerization is sufficient to impair Wnt target gene induction. **A.** Representative images of PLA foci of endogenous β-catenin and endogenous E-cadherin. DAPI was used for nuclear staining. Scale bar, 10 μm. **B.** β-catenin/E-cadherin PLA quantification from experiment in (A). As PLA foci cannot be unambiguously assigned to each cell, they were quantified for each view, and the average number of foci/cell is shown. More than 70 cells were analyzed per experiment. Shown is mean ± SEM from 3 independent experiments. Welch’s *t*-test. **C.** Representative images of smRNA-FISH of endogenous *CTNNB1* and endogenous *CDH1* in iPSCs. Scale bar, 10 μm. **D.** Quantification of (C). Percent of *CDH1* mRNAs co-localized with endogenous *CTNNB1* is shown as mean ± SEM from 2 independent experiments. More than 50 cells were analyzed per experiment. Welch’s *t*-test. **E.** Quantification of β-catenin/BCL9L PLA foci at DE 4h. Cells analyzed, Ctrl, *N* = 152; Udel, *N* = 165. Shown is mean ± SEM from 2 independent experiments. Welch’s *t*-test. ****, *P* = 9 x 10^−28^. **F.** β-catenin RNA-IP from Fig. 4M, followed by qRT-PCR for *BCL9L* mRNA, shown as mean ± SD from 2 independent experiments. Welch’s *t*-test. **G.** β-catenin/TCF7L2 PLA time course. Shown is mean ± SEM from 2 independent experiments. 0h, Ctrl, *N* = 162, Udel, *N* = 153; 2h, Ctrl, *N* = 163, Udel, *N* = 162; 4h, *N* = 81, Udel, *N* = 91; 6h, *N* = 160, Udel, *N* = 127. Welch’s *t*-test; ****, *P* < 0.0001. **H.** As in (G), but β-catenin/LEF1 PLA is shown. 0h, Ctrl, *N* =170, Udel, *N* = 168; 2h, Ctrl, *N* = 182, Udel, *N* = 149; 4h, *N* = 144, Udel, *N* = 161; 6h, *N* = 171, Udel, *N* = 168; Welch’s *t*-test; ****, *P* < 0.0001. **I.** QRT-PCR of *LEF1* mRNA at the indicated timepoints, shown as in Fig. 2B and performed in parallel to PLA time course from (H). **J.** As in (I), but shown is *TCF7L2* mRNA. **K.** RNA bifold prediction of mRNA heterodimerization of the *CTNNB1* and *LEF1* 3′UTRs, which occurs in the *CTNNB1* dimerization site 1. This region is also the binding site of the *TCF7L2* 3′UTR. Shown as in Fig. 5D. **L.** Intermolecular RNA base pairing (bp) strength in the *CTNNB1* 3′UTR dimerization site 1, when bound to the *TCF7L2* or *LEF1* 3′UTRs. Wilcoxon test, *P* = 0.001. **M.** Schematic of sequential generation of co-folded β-catenin complexes, whose formation is initiated at the mRNA level through intermolecular 3′UTR-3′UTR interactions. **N.** Schematic of the *CTNNB1-TCF7L1* and *CTNNB1-TCF7L2* 3′UTR-3′UTR interactions. Addition of six ASOs blocks both *CTNNB1* 3′UTR dimerization sites. **O.** *LEF1*, *AXIN2, TBXT* mRNA FC at DE 4h in WT iPSCs transfected with three Ctrl ASOs compared to 6-ASOs blocking the *CTNNB1* 3′UTR, as shown in (N). *GAPDH* was used for normalization. Mock, RNAiMAX only. Shown is mean ± SD from 3 independent experiments. Welch’s *t*-test; *, *P* < 0.05; ***, *P* < 0.001. **P.** Scatter plot showing differentially expressed genes as log2FC in *CTNNB1* Udel versus Ctrl (x-axis) and log2FC in 6-ASO treated versus Ctrl ASO treated (y-axis) cells at DE 4h. Genes with significant changes (log2|FC| > 0.58) in either group are shown in blue (*N* = 114), together with the Pearson’s correlation coefficient R = 0.75, *P* = 2.0 x 10^−21^. Dashed lines indicate a FC of 1.5. Selected genes are indicated.

The 3′UTR independence of the protein interaction is further supported by functional experiments. Previous reports suggested that β-catenin promotes E-cadherin localization to the plasma membrane^73^, a finding that we confirmed with *CTNNB1* KO cells (Fig. S7B, S7C). However, in *CTNNB1* Udel cells, where β-catenin protein is present at normal levels, E-cadherin cell surface expression was not diminished (Fig. S7B, S7C). Taken together, β-catenin protein is required for E-cadherin trafficking and cell adhesion, but this protein-protein interaction is not pre-established at the mRNA level.

### Increased co-activator co-localization with β-catenin co-folded complexes

β-catenin also binds to a large number of co-activators, which allow full Wnt target gene induction^23,24,30^. We tested whether β-catenin interaction with either p300 or BCL9L protein requires the *CTNNB1* 3′UTR. Out of all reported β-catenin-interacting co-activators^24^, the two proteins were tested, because they are essential for DE differentiation^34^, suggesting that they could be important in the investigated cellular context. We observed a significant reduction in either β-catenin/BCL9L or β-catenin/p300 protein co-localization in *CTNNB1* Udel cells by PLA (Fig. 6E, S7D).

However, according to β-catenin RNA-IP, there was no evidence for protein co-folding (Fig. 6F, S7E), suggesting that increased β-catenin/co-activator binding is not caused by co-translational co-folding and likely happens in the nucleus. We speculate that the co-folded β-catenin-TCF7L2 complex could have a higher valency, thus getting better retained at the chromatin. It is also possible that the co-folded complex has a new binding surface for co-activators or that the co-activator binding sites in β-catenin are better accessible in the context of the co-folded complex.

### LEF1 displaces TCF7L2 from its co-folded complex with β-catenin

β-catenin binds to all four Wnt transcription factors^24,29^. During the switch from repression to Wnt activation in the course of DE differentiation, binding of TCF7L1 and TCF7L2 to β-catenin turns these initially repressing factors into activators, which will eventually be replaced by LEF1 and TCF7 (Fig. 3H)^29,74^. Although LEF1 was reported to assemble co-translationally with β-catenin^61^, 3′UTR dependence cannot be studied directly in our experimental system, because mRNA expression of *LEF1* is strongly reduced in *CTNNB1* Udel cells (Fig. 2B).

We performed a PLA time course to investigate the dynamics of β-catenin/Wnt transcription factor co-localization during early timepoints after DE induction. In Ctrl cells, β-catenin/TCF7L2 interaction was highest at 4h, followed by a sharp drop at the 6h timepoint (Fig. 6G). In contrast, at the 6h timepoint, β-catenin/LEF1 co-localization was highest within the time frame of the experiment, but was barely detectable in the first four hours, as *LEF1* mRNA and protein first need to be synthesized (Fig. 3G, 6H, 6I). Together these data indicate that LEF1 displaces TCF7L2 in the co-folded complex with β-catenin. Efficient protein exchange requires the *CTNNB1* 3′UTR, as both the sharp drop in β-catenin/TCF7L2 and the rapid increase in β-catenin/LEF1 are blunted in Udel cells (Fig. 6G, 6H). In Udel cells, β-catenin/LEF1 co-localization is reduced, and β-catenin/TCF7L2 co-localization persists beyond the 4h timepoint.

With this experimental system we cannot clearly distinguish between a 3′UTR-dependent or a LEF1 protein abundance-dependent effect in displacing TCF7L2 from the complex with β-catenin at the 6h timepoint. Nevertheless, the corresponding mRNA levels, which were measured in parallel, do not explain the observed pattern in β-catenin/Wnt transcription factor complexes (Fig. 6I, 6J).

RNA bifold predicted a 169-nt long intermolecular interaction between the 3′UTRs of *CTNNB1* and *LEF1*, where 95 nt base-pair intermolecularly (Fig. 6K, Table S5). The mRNA heterodimerization occurs within the *TCF7L2* binding site on the *CTNNB1* 3′UTR (Fig. 6K).

Intriguingly, base pairing of the *CTNNB1-LEF1* RNA dimer is predicted to be significantly stronger than base pairing of the *CTNNB1-TCF7L2* RNA dimer (Fig. 6L). Based on these results, we propose the following model (Fig. 6M). Four to 6h after DE induction, enough *LEF1* mRNA has been synthesized (Fig. 6I). More *LEF1* mRNA together with a higher RNA binding affinity between *CTNNB1* and *LEF1* enables the *LEF1* 3′UTR to displace *TCF7L2* from the *CTNNB1* mRNA heterodimer. In this model, rapid exchange of protein binding partners, which allows efficient sequential generation of co-folded β-catenin complexes, is initiated at the mRNA level through intermolecular 3′UTR-3′UTR interactions.

Interestingly, a similar scenario is predicted for displacement of *TCF7L1* by the *TCF7* 3′UTR. The *CTNNB1-TCF7* dimer is predicted to have a length of 61 nt with 52 of them involved in intermolecular base pairing (Fig. S7F). It occurs in the *TCF7L1* binding site on the *CTNNB1* 3′UTR. The base pairing of the *CTNNB1-TCF7* RNA dimer is predicted to be significantly stronger than the *CTNNB1-TCF7L1* RNA dimer (Fig. S7G); therefore, increasing levels of *TCF7* mRNA, induced through Wnt activation (Fig. 3E), likely replaces *TCF7L1* in the *CTNNB1* 3′UTR dimer.

### The *CTNNB1* 3′UTR heterodimers are necessary to drive the Wnt transcriptional program

The two *CTNNB1* 3′UTR dimerization sites are crucial for formation of β-catenin/Wnt transcription factor complexes, but are they sufficient for induction of the Wnt transcription program? We investigated whether blocking both *CTNNB1* dimerization sites with six ASOs (ASOs 12-14 and 22-24, termed 6-ASOs) impairs Wnt target gene induction (Fig. 6N). Addition of 6-ASOs did not change β-catenin protein levels but significantly reduced induction of several Wnt target genes, when compared with Ctrl ASOs (Fig. 6O, S7H). Although the ASOs do not degrade the mRNA, it is currently unclear whether they just block the sequence or also alter the 3′UTR structure. The 6-ASO-treated cells also reduced expression of Wnt targets in a colon cancer cell line (Fig. S7I).

To examine if all Wnt target genes are affected by the 6-ASO treatment, we performed RNA-seq at DE 4h. We compared the fold-change in gene expression in Udel versus Ctrl and in 6-ASO-treated versus Ctrl ASO cells (Fig. 6P, Table S4). We observed a strong correlation between the gene expression changes upon *CTNNB1* 3′UTR deletion and 6-ASO-mediated blocking of both *CTNNB1* 3′UTR heterodimers (Fig. 6P). This experiment demonstrates that formation of 3′UTR heterodimers is necessary for full Wnt target gene induction during iPSC differentiation. This result indicates that post-translational protein-protein interactions between β-catenin and the Wnt transcription factors TCF7L1 or TCF7L2 are insufficient for rapid Wnt target gene induction.

### C-clamp-regulated Wnt targets are strongly reduced upon *CTNNB1* 3′UTR deletion

Next, we set out to learn the differences between the co-folded and post-translationally formed Wnt transcription factor complexes. Examining the differentially expressed genes at DE 4h revealed that not all Wnt targets are equally affected in *CTNNB1* Udel versus Ctrl cells (Fig. 7A). The strongest reduction in expression was observed for *SP5, TBXT, NKD1*, and *LEF1*, which are known as C-clamp-regulated genes (Table S4)^29,75,76^.

**Figure 7.**
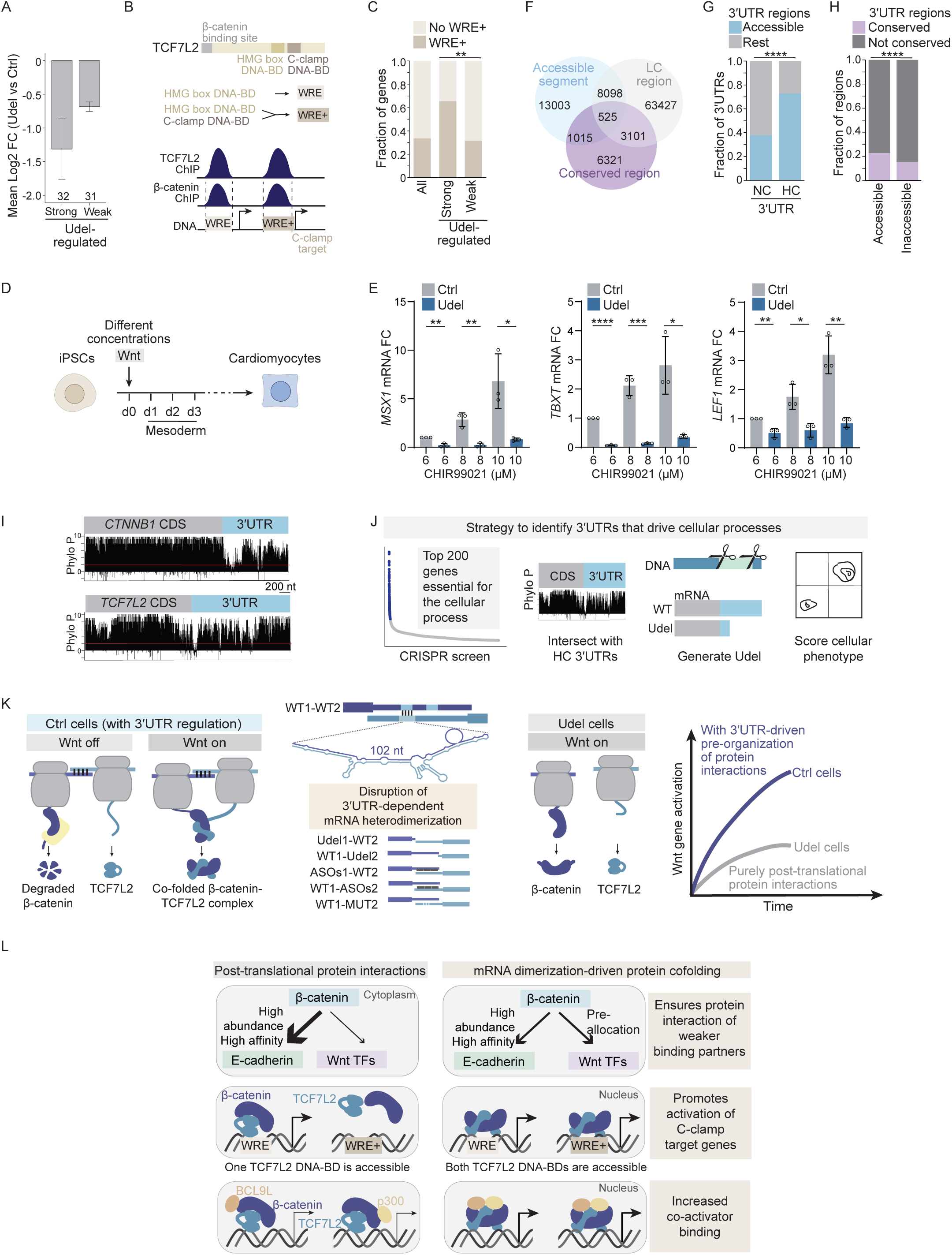
3′UTR-driven co-folded protein complexes provide quantitative and qualitative advantages over post-translationally formed interactions. **A.** Target genes strongly or weakly affected in *CTNNB1* Udel cells at DE 4h, obtained by median split. Mann Whitney test, *P* = 8.3 x 10^−6^. **B.** Schematic for genome-wide identification of C-clamp target genes. Protein model of TCF7L2 with protein domains is shown. DNA-BD, DNA-binding domain. The HMG box DNA-BD binds to the WRE, whereas combined binding of the HMG box and C-c lamp DNA-BD to the WRE+ is required for C-clamp target gene induction. Occurrence of the WRE+ was determined in genomic regions co-bound by TCF7L2 and β-catenin. **C.** The WRE+ is significantly enriched among genes strongly affected in Udel cells. X^2^ = 11.9; *P* = 0.0006. All, all genes expressed in Ctrl or Udel cells at DE 4h (*N* = 10,405). **D.** Schematic of iPSC to mesoderm and cardiomyocyte differentiation. **E.** mRNA expression of indicated C-clamp targets at 48h post-Wnt induction, obtained with increasing amounts of CHIR99021 and normalized to *GAPDH*. Shown is mean ± SD from 3 independent experiments. *T*-test; *, *P* < 0.05; **, *P* < 0.01. **F.** Venn diagram showing the relationships between accessible regions, LC regions, and conserved regions across human 3′UTRs. **G.** Accessible 3′UTR regions are significantly enriched among HC 3′UTRs. HC 3′UTRs with accessible regions, *N* = 1661; HC 3′UTRs without accessible regions, *N* = 621; NC 3′UTRs with accessible regions, *N* = 2530; NC 3′UTRs without accessible regions, *N* = 4132. Mann Whitney test, *P* = 7 x 10^−182^. **H.** Conserved 3′UTR regions are significantly enriched within accessible compared to inaccessible 3′UTR segments. Accessible region is conserved, *N* = 1382; accessible region is not conserved, *N* = 4693; inaccessible region is conserved, *N* = 5766; inaccessible region is not conserved, *N* = 32112. X^2^ = 219; *P* < 0.0001. **I.** PhyloP scores of CDS and 3′UTR of *CTNNB1* and *TCF7L2* mRNAs. **J.** Schematic representation of strategy to identify 3′UTR with the potential to drive cellular processes. **K.** Model for mRNA heterodimerization-driven protein co-folding, necessary for rapid and high Wnt target gene induction.\ **L.** Model, showing quantitative and qualitative advantages of co-folded β-catenin/TCF7L2 protein complexes over post-translationally formed protein-protein interactions.

All Wnt transcription factors contain a high-mobility group (HMG) box, which serves as DNA-binding domain for the Wnt-response element (WRE). Certain TCF7L2 splice isoforms, which are expressed in our system^77^, contain an additional, cysteine-rich DNA-binding domain, called C-clamp (Fig. 7B). It binds to a GC-rich helper sequence in the vicinity of the WRE and changes the set of induced Wnt targets^29,77,78^.

We investigated whether expression of C-clamp targets was preferentially reduced in *CTNNB1* Udel cells. We analyzed ChIP-seq data from human ESCs and focused on peaks co-bound by TCF7L2 and β-catenin 4h post-Wnt induction^27,79^. We identified the peaks containing the WRE plus helper sequence (WRE+) and identified their closest genes (Fig. 7B, Table S6). The genes significantly reduced in Udel vs Ctrl cells were split into two groups, containing strongly or weakly reduced target genes (Fig. 7A). Among the genes strongly reduced in Udel cells, the WRE+ peaks were significantly overrepresented (Fig. 7C). However, no enrichment of WRE+ sequences was detected among weakly reduced genes (Fig. 7C). Our results suggest that the C-clamp DNA-binding domain is more active when TCF7L2 is part of the co-folded complex, whereas it is less active when TCF7L2 protein is synthesized individually.

We tested C-clamp gene induction in another iPSC differentiation pathway. We observed that in *CTNNB1* Udel cells, the generation of cardiomyocytes was strongly impaired (Fig. S7J). Formation of mesoderm is the first step of iPSC-induced cardiomyocyte differentiation and requires high levels of Wnt signaling (Fig. 7D)^50,51,80^. We examined gene induction at different levels of Wnt activation and observed a correlation between Wnt signaling strength and gene induction in Ctrl cells (Fig. 7E). In *CTNNB1* Udel cells however, target gene induction was strongly defective and could not be overcome with higher CHIR99021 concentrations (Fig. 7E, S7K). Higher CHIR99021 concentrations did not affect β-catenin protein levels in this context (Fig. S7L). Our results show that the *CTNNB1* 3′UTR is especially important for C-clamp target gene activation in pathways that require high Wnt activation. Taken together, these observations indicate a qualitative difference between co-folded β-catenin/Wnt transcription factor complexes and post-translationally formed interactions.

### HC 3′UTRs are enriched for sequence and structural features that promote mRNA heterodimerization

Lastly, we set out to estimate how widespread mRNA heterodimerization could be across human 3′UTRs. The *CTNNB1* mRNA heterodimerization region is characterized by three features: it is ‘intrinsically unpaired’, contains ‘low complexity’ (LC) regions, and its sequence is conserved (Fig. 5D, 5G, 5K). We set out to identify regions with similar sequence and structural characteristics across all 3′UTRs. Intrinsically unpaired 3′UTR regions have weak RNA self-interaction probability, and therefore, can be considered as accessible regions. We used RNAfold to determine RNA self-interaction probability for each nt across individual 3′UTRs. We counted 22,264 accessible 3′UTR regions that have a minimum length of 75 nt and a self-base pairing probability smaller than 0.3 (Fig. 7F, Table S7).

With respect to the sequence features of 3′UTR heterodimerization regions, we observed that all so far known regions contain LC regions. For example, the *CTNNB1-TCF7L1* interaction site contains a 25-nt-long U-stretch that base pairs with As, whereas the *CTNNB1-TCF7L2* interaction site contains a 26-nt-long region, where 58% of the nucleotides are Gs that pair with Cs in the interacting RNA (Fig. 5D, 5G). Although the LC regions only represent 20-25% of the entire dimerization regions, they could act as nucleation sites for the 3′UTR-3′UTR interactions. RNA regions containing repeats were previously found to be overrepresented among RNA-RNA interaction sites^81^. Here, we identified all LC regions in human 3′UTRs with a minimum length of 45 nts and with overrepresentation of one nucleotide. We observed 8,623 accessible 3′UTR segments that contain LC regions (Fig. 7F, Table S7).

The *CTNNB1* mRNA heterodimerization region has a high phyloP sequence conservation score (Fig. 5K). We identified all accessible 3′UTR segments with high sequence conservation. When we required that more than half of the nt within an accessible segment have phyloP conservation scores greater than 2, we detected 1,540 conserved accessible 3′UTR segments (Fig. 7F, Table S7). Among them, 525 3′UTR segments are accessible, contain LC regions, and are conserved. Our analysis indicates that hundreds of 3′UTR regions have features that mimic the *CTNNB1* heterodimerization domain. Therefore, our results strongly suggest that 3′UTRs beyond the β-catenin/Wnt transcription system use 3′UTR-driven protein co-folding.

Based on our findings so far, we propose that 3′UTRs need to have accessible regions to form intermolecular interactions. Among the tested *CTNNB1* 3′UTR heterodimers, we could not verify interactions with *CDH1*, *EP300* or *BCL9L* (Fig. 6A-F, S7E). Importantly, their predicted heterodimerization sites map to inaccessible 3′UTR regions, whereas the validated 3′UTR heterodimers map to accessible regions (Fig. S7M). We propose that accessible RNA regions are necessary for mRNA heterodimerization, whereas LC regions and conserved nt may play promoting roles.

We asked whether accessible regions are overrepresented within a specific 3′UTR subgroup and tested whether they are enriched among HC 3′UTRs. We applied previously used criteria to identify HC and non-conserved (NC) 3′UTRs^9^. Among HC 3′UTRs, 73% have accessible regions, whereas they are only present in 38% of NC 3′UTRs (Fig. 7G). This result suggests that HC 3′UTRs have a higher propensity to form intermolecular 3′UTR interactions.

We wondered whether accessible 3′UTR regions are under evolutionary pressure. We compared the distribution of conserved 3′UTR regions among accessible and inaccessible 3′UTR segments. We observed that 23% of accessible 3′UTR segments are conserved, whereas only 15% of inaccessible regions are considered conserved (Fig. 7H), indicating that accessible regions are under evolutionary pressure to be maintained.

### Strategy to identify HC 3′UTRs that potentially drive cellular processes

Here, we present a strategy to identify 3′UTRs that potentially drive cellular processes. In the *CTNNB1* and *TCF7L2* mRNAs, the CDS and 3′UTR phyloP conservation scores are similar, suggesting that these HC 3′UTRs regulate crucial features of the mRNA-encoded proteins (Fig. 7I). Although the high 3′UTR sequence conservation scores imply functional importance, the biological context and the mechanism by which these 3′UTRs regulate cellular processes had been unknown.

To identify functionally relevant 3′UTRs within a specific biological context, we selected a cell state change in which the positive and negative regulators had previously been identified with a genome-wide CRISPR screen^34^. We intersected the essential genes in this pathway with HC 3′UTRs and generated genomic 3′UTR deletions. We chose a phenotypic readout to identify potential 3′UTR effects, as the consequences of 3′UTR deletions were unknown. With this strategy (Fig. 7J), most tested HC 3′UTRs had regulatory roles and a subset, such as the *CTNNB1* 3′UTR, even acts as driver of a specific cellular process. Our strategy can be applied to any cellular process, where the positive and negative regulators are known.

### Rapid Wnt target gene induction requires 3′UTR-driven protein co-folding

Investigating the mechanism by which the *CTNNB1* 3′UTR regulates DE differentiation, we discovered the existence of mRNA heterodimers that are translated into co-folded protein dimers (Fig. 7K). Based on our experimental evidence, we propose the following model: The heterodimers between *CTNNB1* and Wnt transcription factor mRNAs already exist before Wnt activation and pre-allocate a pool of β-catenin to act in the Wnt pathway (Fig. 4D). In the absence of Wnt ligand, β-catenin protein is destabilized^41–43^, preventing the generation of β-catenin-containing co-folded transcription factor complexes (Fig. S5G, S5H). After Wnt activation, β-catenin nascent chains co-fold with Wnt transcription factors (Fig. 4I-M, S5I, S5J). The 3′UTR-driven pre-allocation ensures that enough β-catenin molecules are available for binding with Wnt transcription factors (Fig. 3K, 4F, 4H). Within co-folded complexes, the E-cadherin binding site within β-catenin is occupied, thus preventing E-cadherin interaction.

Disruption of mRNA heterodimerization was accomplished through genetic 3′UTR deletions, ASO-mediated blocking, or mutation of the 3′UTR dimerization region (Fig. 7K). Disruption of mRNA heterodimerization generates a cellular condition, where only individual β-catenin and Wnt transcription factor molecules are synthesized, therefore, relying entirely on post-translational protein-protein interactions, which are insufficient for rapid and high Wnt target gene induction (Fig. 2B, 2H, 6P, 7E). 3′UTR-mediated protein co-folding is necessary for full Wnt target gene expression in several biological models, including iPSC differentiation to definitive endoderm, mesoderm, and cardiomyocytes as well as during zebrafish embryogenesis (Fig. 1G, 2K, 2L, 7E, S7J).

### Advantages of protein co-folding over post-translational interactions

Formation of *CTNNB1* mRNA heterodimers allows pre-allocation of a pool of β-catenin to function in the Wnt pathway (Fig. 4D, 4F, 4H). This regulatory mode overcomes the strong disadvantage of Wnt transcription factors when competing with E-cadherin for β-catenin binding.

The in vitro binding affinity between E-cadherin and β-catenin is 500-fold higher than between β-catenin and the Wnt transcription factors^82^. Moreover, in iPSCs, E-cadherin protein is expressed at 12-56-fold higher levels than the Wnt transcription factors TCF7L1 and TCF7L2^83^, indicating that post-translational β-catenin/E-cadherin protein interactions are sufficient for their biological effects (Fig. S7C). In contrast, 3′UTR-dependent mRNA heterodimerization between *CTNNB1* and the Wnt transcription factor mRNAs is necessary to overcome the disadvantage for protein interaction, thus allowing more β-catenin molecules to act in the Wnt pathway (Fig. 7L).

Our findings on preferential regulation of C-clamp targets by the 3′UTR-driven co-folded protein complexes further suggest a qualitative difference between individual TCF7L2 molecules and TCF7L2 co-folded complexes (Fig. 7A-E, 7L). The molecular details of this difference are currently not clear, but it could be that the C-clamp DNA-binding domain is better accessible in the co-folded complex, as the C-clamp is located in an IDR region with misfolding potential^9^. As *LEF1* is one of the strongest C-clamp genes, 3′UTR-driven protein co-folding could be the major mechanism for rapid induction of the Wnt gene expression program.

Lastly, full Wnt target gene expression requires the binding of co-activators to the β-catenin/Wnt transcription factor complexes^23,24,29,30^. We observed increased co-activator recruitment in Ctrl versus *CTNNB1* Udel cells (Fig. 6E, S7D). Although the responsible mechanisms are currently unknown, more β-catenin co-activator binding likely contributes to the stronger Wnt target gene activation observed in the presence of the *CTNNB1* 3′UTR (Fig. 7L).

## Discussion

It has been thought that individual mRNAs are translated into individual protein molecules. Here, we discovered that mRNA heterodimers are translated into co-folded protein dimers. Pre-organizing protein binding events through intermolecular 3′UTR-3′UTR interactions ensures co-localization of proteins. This is in contrast to post-translational protein interactions, where individual molecules eventually meet through random collisions.

We propose that 3′UTR-driven protein co-folding is especially important for cell state changes that happen at short timescales. It allows faster induction of transcriptional programs, which is crucial for development (Fig. 6O, 6P, 7E). For proteins with several binding partners, pre-organization of protein interactions at the mRNA level ensures that the weaker partner is not outcompeted (Fig. 7K, 7L). Moreover, 3′UTR-driven co-folding between a structured and a largely intrinsically disordered protein may allow IDRs to adopt unique and stable conformations during protein synthesis, providing an explanation for the preferential activation of C-clamp target genes by the 3′UTR-dependent complex (Fig. 7L). Our data suggest that 3′UTR-driven mRNA heterodimerization increases the quantity and changes the quality of the generated protein complexes, allowing rapid and high induction of the Wnt transcriptional program during iPSC differentiation and zebrafish embryogenesis (Fig. 7K, 7L).

### mRNA dimerization events pre-arrange future protein interactions

Our most surprising finding was that future protein-protein interactions are pre-established at the level of their encoding mRNAs (Fig. 4D, 4F, 4H, 6G-M). Even before Wnt pathway activation, the protein dimers that will switch the Wnt program from repression to activation are present as mRNA dimers (Fig. 4B-D). As genetic disruption or their blocking through 3′UTR-targeting ASOs is sufficient to impair the transcriptional effects of the protein dimers, our findings reveal that the mRNA dimers are the drivers of the Wnt gene expression program (Fig. 2H, 6P).

Pre-organization of protein interactions at the mRNA level seems to serve two purposes. It allows a pool of β-catenin molecules to be allocated to act in the Wnt pathway. This arrangement allows binding of weaker β-catenin interaction partners and enables their rapid exchange within β-catenin complexes. It seems that sequentially formed β-catenin complexes are established first at the level of mRNA dimerization. Shortly after DE induction, the β-catenin/TCF7L2 dimer forms, which is later replaced by the β-catenin/LEF1 dimer, a process that is strongly promoted by the *CTNNB1* 3′UTR (Fig. 6G, 6H). Increasing amounts of *LEF1* mRNA paired with its stronger RNA binding affinity to the *CTNNB1* 3′UTR displace *TCF7L2*, thus promoting protein exchange of the Wnt transcription factors within the β-catenin co-folded complexes (Fig. 6M).

### HC 3′UTRs allow mRNAs to play active roles during protein synthesis

mRNAs are regarded as templates for protein synthesis. This traditional viewpoint assigns passive roles to mRNAs and allows 3′UTRs to have fine-tuning roles. However, the existence of mRNAs whose 3′UTRs have similar conservation scores as their CDS (Fig. 7I), indicated that these 3′UTRs have essential biological roles. However, it had been unclear in what cellular contexts individual HC 3′UTRs would realize their regulatory roles. Here, we developed a strategy to identify cellular contexts where specific 3′UTRs may act as drivers of cellular pathways (Fig. 7J).

Through genetic disruption of specific 3′UTR regions without altering the sequence and abundance of endogenous proteins, our approach revealed the existence of mRNA-driven protein organization. Moreover, we propose that deletion of endogenous 3′UTRs can be used as strategy to test whether post-translational protein-protein interactions are sufficient for the regulation of a specific pathway. As more than 2,700 mRNAs have HC 3′UTRs, many of which have regions with a propensity to form mRNA heterodimers (Fig. 7F), it is likely that several signaling-induced transcriptional programs will rely on the presence of mRNA 3′UTRs. It will be important to identify additional 3′UTRs that act as drivers of cellular processes to gauge how widespread this regulatory mechanism is. We hypothesize that the blocking of specific 3′UTR regions could represent a new strategy for the modulation of disease programs in the future (Fig. S7I).

Co-translational protein complex assembly has been widely observed^60,61^. In most reported cases, two structured protein domains co-translationally interact^60,61,64,84–87^. The co-folded complexes described by us contain a predominantly structured protein, β-catenin, and one of the Wnt transcription factors, which are predicted to be nearly entirely disordered (Fig. S7N). We speculate that mRNA dimerization-induced proximity provides a mechanism by which the nascent chains of intrinsically disordered proteins use their binding partners as scaffolds for co-translational protein folding.

As a result, an IDR-containing protein can be present in two distinct stable conformations within the same cell type. One conformation is adopted when the individual protein is synthesized, whereas the other conformation occurs in the context of the co-folded complex. Based on our previous results^9^, only small peptides in the IDR participate in protein folding, thus adopting stable, irreversible conformations, whereas more than 90% of IDR sequence is still present as rapidly interconverting conformations. This scenario is consistent with our observation that TCF7L2 can regulate WRE- and WRE+ containing Wnt target genes.

### 3′UTR regions have evolved features that promote intermolecular RNA interactions

RNAs with regions of weak self-structure have a strong potential for interaction with other RNAs^88,89^. We call regions of weak RNA self-structure, accessible regions, and found that a substantial proportion is conserved. Importantly, conserved 3′UTR regions are overrepresented among accessible versus inaccessible 3′UTR segments (Fig. 7H), indicating that 3′UTRs preserve their potential to form intermolecular RNA interactions.

Moreover, we observed that accessible 3′UTR regions are concentrated within HC 3′UTRs (Fig. 7G). We previously observed that mRNAs with HC 3′UTRs are strongly enriched within mRNA-scaffolded meshlike condensates^9^. Moreover, engineered intermolecular RNA-RNA interactions are capable of generating condensates with meshlike morphology^88^. Taken together, these findings suggest that 3′UTR-driven mRNA heterodimerization events could serve as the endogenous scaffolds of meshlike cytoplasmic condensates^3,5,15,90^. Moreover, co-localization of HC 3′UTRs within cytoplasmic meshlike condensates may represent a mechanism by which two different mRNAs meet to form mRNA dimers. However, these hypotheses need experimental testing in the future.

### How mRNA dimerization is regulated is currently unknown

mRNA heterodimerization events, including 3′UTR-3′UTR interactions have been observed previously in psoralen-based chemical crosslinking datasets^81^. Moreover, heterodimerization between lincRNAs and mRNAs were previously observed in Alu repeats within 3′UTRs, where binding of the double-stranded RNA-binding protein Stauffen causes mRNA destabilization^91^. As 3′UTRs are strongly bound by a large number of RNA-binding proteins, we suspect that different RNA-binding proteins promote or inhibit formation of specific mRNA dimers. Identifying the positive and negative regulators of mRNA dimerization will be important to manipulate 3′UTR-driven pathways in the future.

## Limitations of the study

Although our computational analysis predicts hundreds of human 3′UTRs with sequence and structural features similar to the *CTNNB1* mRNA heterodimerization region, we have not shown that 3′UTR-dependent protein co-folding occurs beyond the β-catenin/Wnt transcription factor system.

Although we have direct evidence for co-translational assembly between β-catenin and the Wnt transcription factors TCF7L1 and TCF7L2, we currently have no insights into the stoichiometry of the heteromeric assemblies. We also do not have structural evidence that the protein conformation of TCF7L2 is different in the co-folded versus the individual protein.

## Supplementary Tables

Table S1. mRNA features of 3′UTR deletion candidates, related to Figure 1, 2. Table S2. Amplicon sequencing, related to Figure 1.

Table S3. ASOs, oligonucleotides, guide RNAs, and probes, related to Key Resources Table.

Table S4. RNA sequencing of Ctrl, Udel, and *CTNNB1* KO clones, related to Figure 2, 6.

Table S5. Features of the experimentally tested mRNA heterodimerization events, related to Figure 5, S7.

Table S6. TCF7L2 and β-catenin ChIP-seq analysis, related to Figure 7.

Table S7. Sequence and structural features across human 3′UTRs, related to Figure 7.

## Acknowledgements

This work was funded by the NIH grant R35GM144046, the Basic Research Innovation Award (BRIA), a grant from the Pershing Square Foundation, William Ackman, and Neri Oxman to C.M. as well as the MSK Core Grant (P30 CA008748). We thank Renhe Luo for initial guidance for DE differentiation, Xiaoping Bao for valuable suggestions on knock-in experiments, and Riley Ogrean for help with generating the *KDM6A* 3′UTR deletion cells. We thank all members of the Mayr lab, and Nancy M. Bonini (University of Pennsylvania) for critical reading of the manuscript and helpful discussions.

## Author contributions

T.C. performed all experiments except the injections into zebrafish eggs, which were performed by N.M.C. N.M.C. and R.M.W. designed and interpreted the zebrafish experiments. S.B. performed the computational analyses with input from C.M. T.C. and C.M. conceived the project, designed the experiments, and wrote the manuscript with input from all authors.

## STAR methods

### RESOURCE AVAILABILITY

#### Lead contact

Further information and requests for resources and reagents should be directed to and will be fulfilled by the Lead contact, Christine Mayr.

#### Materials availability

• Plasmids generated in this study will be deposited to Addgene.

• The gene-edited iPSC and ESC clonal lines generated in this study are available from the Lead contact with a completed Materials Transfer Agreement.

#### Data and code availability

Processed RNA-seq data are reported in Table S4, and the raw data are accessible through GEO accession number GSE329836. C-clamp targets were identified from publicly available ChIP-seq data (GSM1505791, GSM1579346) and are reported in Table S6. Sequence and structural features of human 3′UTRs are reported in Table S7. Analysis code for C-clamp target identification, accessible, LC, and conserved 3′UTR regions were deposited at Zenodo (https://zenodo.org/records/24375870). Any additional information required to reanalyze the data reported in this paper is available from the Lead contact upon request.

### Experimental model

#### Cell lines

This study used two established human pluripotent stem cell lines were used. The iCas9 *SOX17^eGFP/+^*HUES8 ESC line was generated by the laboratory of Danwei Huangfu^34^. The iCas9 731.2B iPSC line was obtained from the Stem Cell core facility (MSKCC)^92^. Both cell lines contain inducible Cas9 inserted at the AAVS1 locus. Undifferentiated iPSCs were maintained in StemFlex medium (Gibco, A3349401) on Matrigel (Corning, 354277) coated plates. ESCs were maintained in E8 medium (Gibco, A1517001) on plates coated with human recombinant vitronectin (Gibco, A14700). All ESC/iPSCs were maintained at 37°C with 5% CO2. Both stem cell lines were dissociated with 0.5 mM EDTA in PBS and passaged every 2–3 days. For single-cell dissociation steps, cells were treated with 10 µM ROCK inhibitor (Stemcell, Y-27632) for 24h after plating. Cells were routinely confirmed to be mycoplasma-free by PCR. All experiments were approved by the Tri-Institutional stem cell initiative ESC research oversight committee (ESCRO protocol #2024-01-003).

#### Generation of 3′UTR deletion ESC or iPSC clones

3′UTR deletion stem cell lines were generated using a pair of guide RNA. Guide RNAs were designed to avoid disruption of the coding sequence or distal mRNA cleavage and polyadenylation signals^93^. Three pairs of guide RNAs were designed using CRISPOR^94^ and were tested in HEK293T cells to select for the highest editing efficiency (data not shown). The guide RNA pairs with the highest editing efficiency were purchased from Synthego (Table S3). Early passage ESC/iPSC were used for editing. To induce Cas9 expression, 1 µg/ml doxycycline was added to the medium 24h before transfection. Cells were dissociated with TrypLE Select Enzyme (Gibco, 12563029) and resuspended in E8 medium supplemented with ROCK inhibitor for both cell lines (we found that maintaining cells in E8 medium during transfection improved editing efficiency compared with StemFlex). Ten nM of each guide RNA was reverse transfected with 2.5 x 10^5^ cells using Lipofectamine RNAiMAX (Invitrogen, 13778075) in a 12-well plate.

Two days after transfection, cells were dissociated into single cells using Accutase (Innovative Cell Technologies, #AT-104). Cells were filtered through a 40 µm cell strainer, and ∼4,000 cells were plated as single cell suspension in a 10 cm plate with ROCK inhibitor-supplemented culture medium. The medium was refreshed daily for 6–7 days.

Single colonies were picked into a 96-well plate and allowed to grow for two days. Cells were then split into two 96-well plates with EDTA: 20% of the cells were plated for maintenance and 80% for genotyping. Cells were collected and lysed with QuickExtract DNA Extraction Solution (Biosearch Technology). PCR was performed to screen for potential Udel clones. Potential Udel clones were expanded, and RT-qPCR was performed for further validation using primers targeting the coding sequence and 3′UTR, respectively. In addition, DNA was collected for Amplicon sequencing. Ultimately, three homozygous independent clones were selected for downstream analysis. Karyotyping was not performed.

#### Generation of *CTNNB1* knockout cell lines

*CTNNB1* knockout (KO) cells were generated in iCas9 731.2B, where a coding sequence-targeting guide RNA (Table S3) was transfected into cells as described above and used for all experiments except the rescue experiment reported in Fig. 3A-D. For the rescue experiment, *CTNNB1* KO cells were generated in 731.2B cells that lack Cas9. In these cells, the coding sequence-targeting guide RNA (Synthego) and S.p. HiFi Cas9 Nuclease V3 (IDT, 1081060) were transfected using Lipofectamine™ Stem (Invitrogen, STEM00008) in E8 medium according to the manufacturer’s instructions. Three homozygous clones were selected by western blot for both KO cell lines for downstream analysis. Karyotyping was not performed.

#### Generation of AAVS1 locus knock-in iPSCs

The AAVS1 knock-in donor plasmid was designed to replace Cas9 in the original vector (Addgene #73500) with the human *CTNNB1* coding sequence and full-length 3′UTR (using the MANE annotated mRNA transcript, Fig. S6A). The plasmid expressing the AAVS1 T2 gRNA in combination with FLAGless eSpCas9(1.1) was obtained from Addgene (#79888). Plasmids were mixed at a 4:1 ratio and transfected using Lipofectamine™ Stem into the 731.2B *CTNNB1* KO cells, described above. At 48h post-transfection, 1 µg/mL puromycin was applied for 10 days of selection.

#### Other cell lines

HEK293T and HCT116 cells were obtained from ATCC and were cultured in high-glucose DMEM supplemented with 10% FBS and 1% penicillin-streptomycin at 37 °C in 5% CO_2_.

#### Generation of *CTNNB1* KO HEK293T and 3′UTR-mutated stable cell lines

HEK293T cells were transfected with Cas9-sgRNA RNP targeting *CTNNB1* as described above. Single-cell clones were isolated and screened by western blot for complete loss of β-catenin. To generate lines expressing *CTNNB1* CDS with mutations in its 3′UTR, *CTNNB1* CDS was fused to the full-length wild-type or mutant 3′UTR and cloned into pcDNA3.1-puro^3^. The mutant 3′UTR was generated by site-direct mutagenesis, replacing the wild-type sequence 5′-GAGGGCTCGAGGGGTGGGCT-3′ with 5′-GACCCCTCGACCCCTCCCCT-3′. The introduced ten G-to-C substitutions are reciprocal to the C-to-G substitutions introduced into the TCF7L2 3′UTR (see below), so that predicted base-paring between the two mRNA is disrupted in each single mutant but restored when the two mutants are co-expressed. Constructs were verified by Sanger sequencing and transfected into *CTNNB1* KO cells with Lipofectamine 3000. Stable integrant cells were selected with 2 µg/ml puromycin for three weeks, leading to polyclonal populations carrying randomly integrated transgene copies. β-catenin expression in each line was assessed by RT-qPCR and western blot.

Flag-tagged *TCF7L2* carrying the full-length wild-type or mutant 3′UTR was cloned into pcDNA3.1. The TCF7L2 isoform (Q9NQB0-1) was amplified by PCR from Addgene plasmid #141884. The full-length *TCF7L2* 3′UTR was amplified from genomic DNA and cloned downstream of the coding sequence. The mutant 3′UTR was generated by site-directed mutagenesis, replacing the wild-type sequence of 5′-ATGCCCTACCCTGCGCCCTT-3′ with 5′-ATGGGGTAGGGTGCGGGGTT-3′ and thereby introducing nine C-to-G substitutions. All constructs were verified by Sanger sequencing.

#### Generation of zebrafish embryos with *ctnnb1* KO or *ctnnb1* 3′UTR deletions

**Transgenic zebrafish husbandry.** The wildtype zebrafish strain Tropical 5D (T5D) was used for all experiments^95^. Adult fish were housed at 28.5 °C under standard husbandry conditions, including a 14:10 h light–dark cycle, regulated pH 7.4, and monitored salinity. Animals were fed daily with live brine shrimp followed by Zeigler pellets. All procedures were conducted in accordance with protocols approved by the Memorial Sloan Kettering Cancer Center Institutional Animal Care and Use Committee (IACUC; protocol 12–05–008). Embryos were obtained through natural spawning and maintained at 28.5 °C in E3 medium (5 mM NaCl, 0.17 mM KCl, 0.33 mM CaCl₂, 0.33 mM MgSO₄). Adult experiments used equal numbers of male and female fish. Sex cannot be determined at the embryonic or early larval stages (1–3 days post-fertilization).

**Zebrafish embryo gene editing using CRISPR-Cas9.** CRISPR-Cas9 ribonucleoprotein (RNP) complexes were assembled using the Alt R system (IDT) following the manufacturer’s guidelines with minor modifications. crRNAs were resuspended to a final concentration of 100 µM in Nuclease Free IDTE or Duplex Buffer. Duplexed guide RNA was prepared by mixing equal volumes of 100 µM crRNA (IDT; Table S3) and 100 µM tracrRNA (IDT, #1072532) to generate a 50 µM gRNA duplex, followed by heating at 95°C for 5 min and cooling to room temperature on the benchtop. RNP complexes were generated by combining duplexed gRNA with Cas9 nuclease (IDT #1081058) at a ratio yielding a final Cas9 concentration of approximately 900 ng/µl. For single guide reactions, 1.11 µl of 50 µM gRNA was mixed with 0.9 µl Cas9 and 5.99 µl buffer to a final volume of 8 µl. For two guide deletion reactions, 0.555 µl of each 50 µM gRNA was used along with 0.9 µl Cas9 and 5.99 µl buffer. All RNP mixtures were incubated at 37 °C for 10 min to promote complex formation, after which 2 µl of Phenol Red was added to reach a final reaction volume of 10 µl.

The RNP complexes were microinjected into one cell–stage zebrafish embryos. Adult fish were paired the evening before injections, and embryos were collected within 10 minutes of fertilization to ensure precise staging. Injection needles were pulled from borosilicate glass capillaries and calibrated to deliver consistent volumes. RNP complexes, consisting of duplexed crRNA:tracrRNA and Cas9 nuclease, were assembled immediately before use and kept on ice. Embryos were aligned in agarose-coated injection dishes, and approximately 1 nl of RNP mix was injected into the yolk or directly into the cell at the one cell stage to maximize editing efficiency. Injected embryos were transferred to E3 medium and maintained at 28.5°C for subsequent phenotypic assessment and genotyping, and amplicon sequencing.

### Method details

#### Definitive endoderm (DE) differentiation

DE differentiation was performed as previously described^34,35^. Cells were dissociated into single cells using TrypLE Select Enzyme. For iCas9 731.2B cells, 1.5 x 10^4^ cells were plated in a 24-well plate in Stemflex medium supplemented with ROCK inhibitor. The medium was refreshed 16h after seeding. For iCas9 *SOX17^eGFP/+^* HUES8 cells, 1.2 x 10^4^ cells were plated for differentiation in E8 media and ROCK inhibitor, the medium was refreshed after 18h. Six hours after medium refresh, cells were washed once with PBS and differentiation was induced with S1/2 media as previously described^35^. S1/2 medium was prepared using MCDB131 supplemented with GlutMax, sodium bicarbonate, glucose, and BSA powder. 30 ng/ml Activin A (Abeomics, 32-7586) was supplemented daily, and 5 µM CHIR99021 (TOCRIS, 4423) was added on the first day.

#### Cardiomyocyte differentiation

Cardiomyocyte differentiation was performed as described previously^50^. iPSCs were dissociated into single cells with Accutase and seeded onto Matrigel-coated 12-well plates at 250,000 cells per well. Cells were cultured in StemFlex until approximately 90% confluency. On day 0, differentiation was induced by replacing the medium with RPMI 1640 (Gibco, 11875093) supplemented with B27 minus insulin (Gibco, A1895601) and 10 µM CHIR99021. After 24h, the medium was replaced with fresh RPMI/B27 minus insulin. On day 3, conditioned medium was collected from each well, combined 1:1 with fresh RPMI/B27 minus insulin, and supplemented with IWP2 (Tocris, 3533) to a final concentration of 5 µM. On day 5, the medium was replaced with fresh RPMI/B27 minus insulin. From day 7 onward, the medium was replaced every 3 days with RPMI supplemented with B27 (Gibco, 17504044).

On day 12, cells were dissociated with 0.25% trypsin and stained with a Live/Dead viability die for 30 min in 0.5% BSA/PBS. Cells were then fixed in 4% formaldehyde, permeabilized in 90% methanol, and stained with Alexa Fuor 647-conjugated anti-cardiac Troponin T (cTnT; BD, 565744, 1:500) or the corresponding IgG1-kappa isotype control (BD, 557732; 1:200) in 0.5% BSA, 0.1% Triton X-100/PBS for 1h at room temperature. The fraction of cTnT-positive cells was determined on a CytoFlex Flow Cytometer and analyzed using FCS Express 7.

Day 2 of iPSC to cardiomyocyte differentiation is considered mesendoderm, whose formation requires high Wnt signaling. The cells at d2 are characterized by high *TBXT* expression.

### Flow cytometry

To determine DE differentiation efficiency using SOX17 and CXCR4 markers, cells were dissociated using TrypLE Select Enzyme and resuspended in FACS buffer (5% FBS, 5 mM EDTA in PBS). For iCas9 731.2B cells, cells were stained with CXCR4-APC antibody (Bio-Techne, FAB170A, 1:50) and LIVE/DEAD Violet dye (Invitrogen, L34961, 1:1000) on ice for 30 min. Cells were then fixed with 4% formaldehyde (Thermo Scientific, 28908) at room temperature for 15 min, followed by permeabilization with 0.1% TritonX-100 for 10min. Cells were subsequently stained with SOX17-PE antibody (BD, 561591, 1:5000) on ice for 30 min, followed by flow cytometry analysis. For iCas9 *SOX17^eGFP/+^* HUES8 cells, cells were stained for CXCR4 on ice for 30 min. After washing with FACS buffer, cells were stained with DAPI (0.5 µg/mL) and analyzed using a Fortessa flow cytometer. The flow cytometry gating strategy is shown in Figure S2A.

For β-catenin and GATA6 protein staining, cells were fixed and permeabilized as described above. β-catenin-PE antibody (BioLegend, 862603, 1:500) and GATA6-PE antibody (Cell Signaling Technology, 26452, 1:200) were used for staining. For surface E-cadherin staining, cells were dissociated using TrypLE Select Enzyme and resuspended in 5% FBS (PBS) buffer. E-cadherin-PE antibody (BioLegend, 147304, 1:500) was used for staining on ice for 30 min. All the antibodies were tested using cells that were negative for the target proteins of interest to ensure specificity. All the data analysis and figure generation were performed in FCS Express 7.

### 3′UTR-blocking antisense oligonucleotides (ASOs)

All ASOs used in this study are listed in Table S3. The nt targeted by the CTNNB1 ASOs are shown in Table S3. The used ASOs were uniformly modified with MOE sugars, a phosphorothioate backbone, and 5-methyl cytosine, and purchased from IDT^96^. For transfection in a 24-well plate, ASOs and 0.5 µl Lipofectamine RNAiMax were diluted in Opti-MEM respectively and incubated for 10 min at room temperature. Cells were dissociated with TrypLE, 1.5 x 10^4^ cells were plated with transfection reagent mix in E8 media. DE differentiation was performed as described above. Samples were harvested at the indicated timepoints.

For the ASO tiling across the *CTNNB1* 3′UTR (Fig. S4B), each experiment includes WT iPSCs that were untreated (UT), transfected with RNAiMAX only (mock), three iPSC sample that were transfected with Ctrl ASO 1, 2 or 3, each, and 8–10 CTNNB1 ASOs at a concentration of 25 nM. CTNNB1 ASOs were randomly grouped in each experiment to rule out batch effects. Each experiment was normalized to the average of three Ctrl ASO treatments. A total of 12 runs of experiments were performed. The ASO tiling across the *GATA6* 3′UTR (Fig. S4E), was performed as described for CTNNB1 ASOs, but WT HUES8 cells were used and each ASO was transfected at a concentration of 50 nM. A total of 11 runs of experiments were performed.

For all ASO experiments, the amount of Ctrl ASOs used was always the same as the amount of target ASOs used. For example, if four ASOs were pooled, the concentration of each ASO was 20 nM and the control ASOs were used at 80 nM (Fig. 5F, 5I, S6H, S6I).

### Amplicon sequencing

Genomic DNA was purified with the Tissue & blood kit (Qiagen, 69504). PCR was performed to amplify a 450 bp region flanking the 3′UTR. PCR products were purified with the PCR Purification Kit (Qiagen, 28104). Library preparation and sequencing were carried out by Genwiz through Amp-EZ services. Paired-end reads were merged using PEAR v0.9.6 with default setting^97^. Unique reads were counted with SeqKit^98^.

### RNA extraction, cDNA synthesis and qRT-PCR

Total RNA was extracted using TRIzol and cDNA was synthesis with qScript master mix (Quantabio, 95048) or SuperScript™ IV VILO™ Master Mix (Invitrogen, 11756050) according to the manufacturer’s instructions. qRT-PCR was performed with SYBR green Master mix (Applied Biosystems; A25742) in QuantStudio 6 flex Real Time PCR system (Applied Biosystems). RT-qPCR was performed with internal control *GAPDH*.

### Cell fractionation

Cell fractionation was performed as pervious described^99^. Briefly, 50 x 10^4^ cells were dissociated by TrypLE enzyme. Cell pellets were resuspended in ice-cold 0.1% NP40 lysis buffer (10 mM Tris-HCl, pH 7.5, 150 mM NaCl), incubated on ice for 5 min, layered over 2.5 volumes of a chilled sucrose cushion (24% sucrose in lysis buffer), and centrifuged at 15,000 g for 10 min at 4°C. The supernatant was collected as the cytoplasmic fraction and the pellet represents the nuclear fraction.

### Immunoblotting

Total protein was collected with Laemmli buffer and boiled at 95°C for 10 min. Proteins were resolved by NuPAGE™ Bis-Tris Mini Protein Gels (Invitrogen) and transferred to a nitrocellulose membrane. Membranes were blocked in Intercept® Blocking Buffer (Li-COR) and incubated with primary antibodies overnight at 4°C. After washing with TBST, membranes were incubated with IRDye® secondary antibodies for 1h at room temperature. Signals were detected using Li-Cor Odyssey image system and quantified with ImageJ. Primary antibodies used in this study were: β-catenin antibody (Abcam, ab32572), 1:5000; β-catenin antibody (BD, 610153), 1:1000; TCF7L2 antibody (Cell Signaling Technology, 2569), 1:1000; TCF7L1 antibody (Cell Signaling Technology, 2883), 1:1000; active-β-catenin antibody (Cell Signaling Technology, 8814), 1:1000; LEF1 antibody (Cell Signaling Technology, 2230), 1:1000; Tubulin antibody (Sigma-Aldrich, T9026), 1:2000; E-cadherin antibody (Cell Signaling Technology, 3195), 1:1000; Actin antibody (Sigma-Aldrich, A2066), 1:000; GATA6 antibody (Cell Signaling Technology, 5851), 1:1000; EOMES antibody (Cell Signaling Technology, 81493), 1:1000; SMAD2 antibody (Cell Signaling Technology, 5339), 1:1000, SMAD2/3 antibody (Cell Signaling Technology, 8685) 1:1000; SMAD4 antibody (Cell Signaling Technology, 46535), 1:1000; CBP antibody (Cell Signaling Technology, 7425). The secondary antibodies used were: IRDye 680RD goat anti-mouse (LI-COR, 926-68070), and IRDye 800CW donkey anti-rabbit (LI-COR, 926-32213).

### RNA-seq

For RNA-seq performed at DE 24h, RNA samples were prepared by TRIzol (Invitrogen, 15596018) extraction according to the manufacturer’s guidelines. 500 ng of total RNA with RIN values of 10 underwent poly(A) selection and TruSeq library preparation through Genewiz Standard RNA-seq services. RNA sequencing was performed on the Illumina® NovaSeq™ platform with paired-end 150 bp reads, generating approximately 20 million reads per sample. For RNA-seq performed at DE 4h, samples were lysed using DAN/RNA Shield (Zymo Research, 1100) and submitted to Plasmidsaurus for RNA-sequencing.

**RNA-seq analysis.** Illumina adaptor sequences were first removed from the paired-end reads through the Trimmomatic v0.39 software^100^. The reads were then aligned to the human genome (hg38/GRCh38) with STAR v2.7.1a^101^. Gene expression was quantified for all samples through HOMER v4.11.1^102^.

To focus on direct Wnt-responsive genes, β-catenin ChIP-seq data were used, which were generated in H1 ESCs after 4h of Wnt3A treatment^27^. The processed bed file was downloaded from the Cistrome data browser and annotated by HOMER annotatePeaks tools using default settings. To identify differentially expressed genes, an FPKM cut-off of 2 was used and the union of genes expressed in Ctrl cells at the stem cell stage and at DE24h was used for further analysis. Differentially expressed genes were identified with a minimum log2|FC| > 0.58 and FDR < 0.05 and intersected with the reported Wnt-responsive genes^27^.

### Chromatin immunoprecipitation (ChIP)

ChIP experiments were performed as described previously^74,103^. 4 x 10^6^ iPSCs were plated for DE differentiation in a 10 cm plate. At 4h post-DE induction, cells were double crosslinked with 0.2 mM disuccinimidyl glutarate (DSG, Thermo Scientific, 20593) for 30 min, followed by 1% formaldehyde (Thermo Scientific, 28908) for 20 min at room temperature. Cells were collected and incubated in Buffer A (25 mM HEPES-NaOH, pH 7.5, 10 mM KCl, 0.1% NP-40 and 1.5 mM MgCl_2_) for 10 min. Cells were pelleted and subjected to MNase (NEB, M0247S, 200 gel units per sample) digestion in digestion buffer (15 mM HEPES-NaOH, pH 7.5, 15 mM KCl, 1 mM CaCl_2_, 3 mM MgCl_2_ and 0.1% NP-40) for 10 min at 37°C. Afterwards, cells were collected in lysis buffer (20 mM Tris-HCl, pH 8.0, 100 mM NaCl, 1 mM EDTA, 0.5 mM EGTA, 0.1% sodium deoxycholate, 0.05% N-lauroylsarcosine, 1% Triton X-100) and sonicated on ice using a sonicator equipped with a microtip probe (MISONIX). Sonication was performed at amplitude 2, pulse ON 5 s, pulse OFF 30 s for 3 cycles. Cell debris was removed, and lysate was collected for immunoprecipitation. Two μg β-catenin antibody (Abcam, ab32572) or IgG was incubated with cell extracts overnight, followed by incubation with Protein G Dynabeads (Invitrogen, 10004D) for 1h. Beads were then collected and washed twice with buffer 1 (0.1% SDS, 0.1% deoxycholate, 1% Triton X-100, 0.15 M NaCl, 1 mM EDTA, 0.5 mM EGTA, 20 mM HEPES, pH 7.6), once with buffer 2 (0.1% SDS, 0.1% sodium deoxycholate, 1% Triton X-100, 0.5 M NaCl, 1 mM EDTA, 0.5 mM EGTA, 20 mM HEPES, pH 7.6), once with buffer 3 (0.25 M LiCl, 0.5% sodium deoxycholate, 0.5% NP-40, 1 mM EDTA, 0.5 mM EGTA, 20 mM HEPES, pH 7.6), and twice with buffer 4 (1 mM EDTA, 0.5 mM EGTA, 20 mM HEPES, pH 7.6) for 5 min each at 4°C. DNA was eluted and de-crosslinked overnight at 65°C. DNA was purified by phenol-chloroform extraction.

For TCF7L2 ChIP experiments, cells were fixed with 1% formaldehyde for 10 min at room temperature. Four µl TCF7L2 antibody (Cell Signaling Technology, 2569S) was used per pulldown sample.

### mRNA co-localization detected by single molecule RNA-FISH

Single molecule (sm) RNA Fluorescence in situ hybridization (RNA-FISH) was performed as described previously with the following modifications^5^. Probes were designed with ProbeDealer^104^. A set of 36 primary probes targeting the respective mRNAs were used. For CTNNB1, only probes targeting the CDS were used (Table S3). Cells grown on coverslips were fixed with 4% formaldehyde for 10 min at room temperature, permeabilized in 0.25% Triton X-100 for 10 min, and washed in PBS. Samples were pre-hybridized in 50% formamide/2x SSC for 5 min at room temperature. Primary probes were diluted to 40 nM in hybridization buffer (50% formamide, 2x SSC, 0.1% yeast tRNA, 10% dextran sulfate, 1% murine RNase inhibitor) and hybridized in a humidified chamber at 37°C for 16h. Unbound probes were removed by two washes in wash buffer (2x SSC, 0.1% Tween-20) at 60°C for 15 min each, followed by one wash at RT for 15 min. Fluorescently labeled secondary probes were diluted to 10 nM each in secondary hybridization buffer (20% ethylene carbonate, 2x SSC) and hybridized for 30 min at room temperature in the dark. Samples were washed twice for 5 min each, in secondary hybridization buffer, stained with DAPI and mounted in ProLong Gold. Imaging was performed on a Nikon CSU-W1 SoRa confocal microscope.

Z-stacks acquired at 0.1 µm interval were deconvolved with NIS-Elements (Nikon) using the default settings. smRNA-FISH spots were segmented and their three-dimensional centroid coordinates and volumes extracted using the 3D objects Counter plugin in ImageJ. Euclidean distances between centroids of the two mRNA species were computed; a pair was scored as interacting when the centroid-to-centroid distance was smaller than the sum of the two average radii.

### Proximity ligation assay (PLA)

Cells were plated on a 12 mm glass slide in 24-well plates. PLA was performed with Duolink PLA kit (Sigma-Aldrich, DUO92007, DUO92004 and DUO92002). Briefly, cells were fixed with 4% formaldehyde for 10 min at room temperature. After three washes with PBS, cells were permeabilized with 0.25% Triton X-100. Blocking was performed with blocking buffer for 1h at 37°C. Primary antibodies (β-catenin antibody (mouse) BD, 610153; TCF7L2 antibody, Cell Signaling Technology, 2569; TCF7L1 antibody, Cell Signaling Technology, 2883; β-catenin antibody (Rabbit), Abcam, ab32572; TLE1 antibody, Santa Cruz, sc-137098) were 1:200 diluted in antibody dilution buffer and incubated on a rotor at 4°C overnight. Slides were then washed and subjected to probe incubation 1h at 37°C. Ligation and amplification were subsequently performed at 37°C for 30 min and 1h 40 min, respectively. Finally, slides were mounted with ProLong Gold mounting medium (Invitrogen, P36935) and single panel imaging was carried out with Nikon Sora confocal microscope.

For each sample, 8–10 views were randomly selected and analyzed. Images were processed with ImageJ^105^. Nuclear masks were generated and ‘Find Maxima’ function was used to count PLA signals in each nucleus. More than 70 cells were analyzed per sample.

### RNA immunoprecipitation (RNA-IP) to assess co-translational protein complex assembly

RNA-IP was performed as previously described, with some modifications^64^. 4 x 10^6^ cells were plated per 10 cm plate for DE differentiation as described above. Two hours post-DE induction, cycloheximide or puromycin were added to the media at a final concentration of 100 µg/ml and returned to the 37°C incubator for 5 min or 10 min, respectively. Subsequently, cells were washed twice with ice-cold PBS and scraped in 1 ml lysis buffer (20 mM HEPES KOH pH 7.5, 150 mM KCl, 10 mM MgCl_2_ and 0.1% (vol/vol) NP-40) supplemented with complete EDTA-free protease inhibitor cocktail (Roche), 40 U/ml SUPERase•In RNase Inhibitor (Invitrogen, AM2696), and cycloheximide or puromycin as needed. Lysates were incubated on ice for 10 min and gently pipetted up and down ten-times. Nuclei were removed by centrifugation at 7,000 g for 5 min. 3% of the clarified extract was saved for input. Two µg β-catenin antibody (Abcam, ab32572), 4 µl TCF7L2 antibody (Cell signaling technology, 2569S) or IgG was incubated with the rest of the extracts for 2h at 4°C with end-over-end mixing. Afterwards, 20 µl of Protein G Dynabeads were added and incubated for 1h. Beads were washed with high salt-containing wash buffer (20 mM HEPES-KOH pH 7.5, 300 mM KCl, 10 mM MgCl_2_ and 0.1% (vol/vol) NP-40) once for 5 min at 4°C with end-over-end mixing, followed by a 3x wash with lysis buffer. RNA was eluted in 100 µl Proteinase K buffer (100 mM NaCl, 10 mM TrisCl pH 7.0, 1 mM EDTA, 0.5% SDS) supplemented with 40 μg proteinase K and incubated at 55°C for 30 min. Three volumes of TRIzol LS reagent (Invitrogen, 10296028) were added, and RNA extraction was performed according to the manufacturer’s instructions.

### Co-immunoprecipitation (co-IP) from cytoplasmic extracts

4 x 10^6^ cells were plated per 10 cm plate for DE differentiation as described above. 2h post-DE induction, cells were harvested in 1 ml lysis buffer (20 mM HEPES KOH pH 7.5, 150 mM KCl, 10 mM MgCl_2_ and 0.1% (vol/vol) NP-40) supplemented with complete EDTA-free protease inhibitor cocktail (Roche). Lysates were incubated on ice for 10 min and gently pipetted up and down to remove nuclei, as described above. Triton X-100 was then added to a final concentration of 1%. For co-IP, 2 µg β-catenin antibody (BD, 610153) or IgG was added and incubated for 2h at 4°C with end-over-end mixing. Subsequently, 20 µl of Protein G Dynabeads were added and incubated for an additional hour. Beads were washed four times with lysis buffer containing 1% TritonX-100 and eluted with 2x Laemmli buffer.

### SunTag/MoonTag translation assay and live-cell imaging of HEK293T cells

*CTNNB1* KO HEK293T cells were transduced with lentivirus encoding scFv-GCN4-sfGFP (Addgene #60907) and Nb-gp41-HaloTag (Addgene #128603)^65,66^. Cells expressing intermediate level of sfGFP and high levels of HaloTag-nanobody were isolated by fluorescence-activated cell sorting. The *CTNNB1* CDS and full-length 3′UTR were cloned into the SunTag translation reporter (Addgene # 74928), replacing the KIF18b coding sequence and PP7 3′UTR; the *TCF7L2* CDS and full-length 3′UTR were cloned in the same manner into the MoonTag translation reporter (Addgene # 128604).

Reporter cells were seeded onto Matrigel-coated glass-bottom imaging dishes and allowed to attach overnight, then co-transfected with the SunTag-*CTNNB1* and MoonTag-*TCF7L2* plasmids at a 4:1 ratio using Lipofectamine 3000. Twenty-four hours after transfection, JF646 HaloTag ligand was added to the medium at 1 µM, 30 min before imaging. Fifteen min later, doxycycline was added at 1 µg/ml to induce reporter transcription. Cells were then washed once with PBS and the medium was replaced with FluoroBrite DMEM (Gibco, A1896701) containing 5 µM CHIR99021. Imaging was performed on a Nikon CSU-W1 SoRa spinning-disk confocal system at a frame interval of 30 ms for up to 1.5 min per field.

### Psoralen-based chemical crosslinking

Psoralen-based chemical crosslinking, followed by pulldown was performed as previously described with some modifications^69^. 1.8 x 10^6^ cells were plated per 6 cm dish. Cells were washed with PBS once and 1 ml 200 mM Psoralen/PBS (Selleck Chemical, S4737) was added per dish and incubated at 37°C for 5 min. Cells were treated with 365 nm UV (UV-A) for 20 min on ice in a Spectrolinker UV crosslinker. Afterwards, cells were scraped from the plate and spun down at 600 g for 5 min. Cell pellets were lysed with TRIzol and total RNA was extracted. 15 µg of total RNA was incubated with 100 pM of biotinylated probes in 1 ml fresh hybridization buffer (2 x SSC, 10 % Formamide, 0.1 % Tween-20, 0.05 mg/ml yeast tRNA (Invitrogen, AM7118), 1 mM EDTA) supplemented with 40 U/ml SUPERase•In RNase Inhibitor. As control, biotinylated probes against *TMSB4X*, a highly abundant mRNA in iPSCs were used^69^. The hybridization was performed on end-over-end rotor over night at 37°C. After hybridization, 40 μl of Dynabeads® MyOne™ Streptavidin C1 beads (Invitrogen, 65001) were used to pull out the RNA complexes. The beads were washed 5x with wash buffer (0.1x NaCl and Sodium citrate (SSC), 0.5% SDS) that had been pre-warmed to 37°C. RNA was eluted from the beads by incubating with 40 µg of proteinase K in 100 µl of PK Buffer (100 mM NaCl, 10 mM TrisCl pH 7.0, 1 mM EDTA, 0.5% SDS) at 50°C for 30 min, and boiling at 95°C for 5 minutes. Three volumes of TRIzol LS reagent were added, and RNA extraction was performed according to the manufacturer’s instructions. The recovered RNA was eluted in 10 µl of nuclease free water and irradiated with 254 nm UV for 1 min for reverse crosslinking. The samples were subsequently used for cDNA synthesis and RT-qPCR.

### Zebrafish experiments

**Zebrafish genotyping.** DNA was isolated from single embryos using QuickExtract DNA Extraction Solution (LGC Biosearch Technologies). Embryos at 3 days post fertilization were transferred individually into 0.2 ml tubes, and residual E3 medium was carefully removed with a pipette. Tubes were placed on ice, followed by the addition of 50 µL QuickExtract solution per embryo. Samples were incubated in a thermocycler at 65°C for 15 min, 68°C for 15 min, and 95°C for 10 min. The resulting lysates were briefly vortexed, centrifuged, and 1 µl of the supernatant was used for PCR amplification using Phusion polymerase (Invitrogen, F630).

**Zebrafish phenotypic assessment.** Embryos were inspected daily on a dissecting microscope, and phenotypes were scored. For imaging, fish were anesthetized with Tricaine methanesulfonate (MS-222, Syndel), prepared as a 4 g/l stock solution adjusted to pH 7.0 and diluted to a working concentration of 0.16 mg/ml. Fish were imaged on a Zeiss AxioZoom V16 microscope with Zen 2.1 software.

### Computational analyses mRNA features

The human MANE gene annotations (GRCH38/hg38) were used for all analyses^106^.

<u>PhyloP sequence conservation in 3′UTRs.</u> The genome-wide phylogenetic *P* value (phyloP conservation score) was obtained from the PHAST package for multiple alignments of 99 vertebrate genomes with the human genome as a bigwig file. The genome-wide phyloP score was mapped to the 3′UTRs of the human GTF (genome-build: GRCh38.p14, genome-build-accession: GCA_000001405.29), and the phyloP conservation values for 3′UTRs were obtained. To determine the number of conserved nt in human 3′UTRs, we used a minimum phyloP score ≥ 2 and designated them as conserved, if at least four consecutive nt had phyloP scores above this cutoff. If a given 3′UTR contains more than 150 conserved nt, is it considered highly conserved (HC). Non-conserved (NC) 3′UTRs contain zero conserved nt, given our criteria. The sum of conserved nt for each 3′UTR is listed in Table S7.

<u>Prediction and visualization of intermolecular RNA-RNA interactions.</u> The RNA bimolecular folding (RNA bifold) program was used to predict intermolecular RNA-RNA interactions using default parameters^67^. If more than one interaction was predicted, we selected the longest, uninterrupted RNA-RNA interaction. Intra-and intermolecular 3′UTR interactions were visualized using RNAfold^70^.

<u>Identification of accessible (intrinsically unpaired) and inaccessible 3′UTR regions.</u> For all human 3′UTRs with a length ≤ 6000 nt, we used their full-length sequences and determined the minimum free energy (MFE) structures using a standalone package of RNAfold (version 2.7.2) from the ViennaRNA package^107^. For each nt, we extracted the values for base pairing probability, which represents a measure for RNA self-interaction strength. We define a 3′UTR region as accessible, if each nt within a minimum of 75 nt has a self-base pairing probability < 0.3, while allowing 10% of the values to be greater than the threshold. The median length of all accessible 3′UTR regions is 120 nt. Inaccessible regions were determined as 3′UTR regions outside of accessible regions with a minimum length of 120 nt.

<u>Identification of accessible conserved 3′UTR regions.</u> For each accessible 3′UTR region, we determined whether it is conserved or non-conserved. An accessible region was considered conserved, if >50% of its nt had phyloP scores > 2. The same criteria were applied to identify conserved inaccessible regions. All regions are listed in Table S7.

<u>Identification of accessible 3′UTR segments that contain low-complexity (LC) regions.</u> We used a custom Python script to identify LC regions. Per our definition, an LC region contains at least 45 consecutive nt, with overrepresentation of a single nt. An LC region is counted, if the A or U content is ≥ 0.5 or if the G or C content is ≥ 0.4. The number of LC regions in each 3′UTR is listed in Table S7. If at least 50% of the LC region overlap with an accessible 3′UTR segment, the region is counted as accessible segment with LC region. The same criteria were applied to identify inaccessible segments with LC regions.

The code to determine accessible regions, LC regions, and conserved regions is available at https://zenodo.org/records/24375870.

<u>Predict base pairing strength of mRNA heterodimerization events.</u> First, we used RNA bifold to determine a specific intermolecular RNA–RNA interaction. To assign numerical values to the interaction, we fused the two RNA sequences together and used it as input for RNAfold. We predicted the MFE structure and determined the base pairing probability for each nt.

### Identification of C-clamp target genes

<u>Identification of genes with WRE or WRE+ using ChIP-seq data.</u> We obtained the chromatin occupancy of TCF7L2 from human ESCs (GSM1505791)^79^. We obtained the chromatin occupancy of β-catenin from H1 hESC treated with Wnt-3a for 4 hours (GSM1579346)^27^. BED files containing peak information from the two datasets were used. The TCF7L2 dataset contained 18,900 peaks. The *CTNNB1* dataset contained 22,725,138 peaks, which were reduced using a score-based filtering approach. We kept all peaks with scores greater than the 90th percentile. After filtering, we obtained 84,463 *CTNNB1* peaks, which were used for downstream analysis.

Overlapping genomic peaks were merged using their outer boundaries to get a unified peak set. We used a custom Python script to scan the underlying genomic sequences on both the positive and negative strands for the Wnt response element (WRE; CTTTGWWS or CTTTGWW^76,78^) and the C-clamp helper motif (RCCG^78^). If a WRE was located within 50 nt of the helper motif, it was designated as WRE+.

Each unified peak was assigned to its closest gene using the bedtools closest command from the BEDTools package (v2.31.0). Peaks overlapping a gene body were directly assigned to that gene. For peaks located between genes, the closest gene was determined based on the minimum genomic distance between the outer boundary of the peak and the corresponding gene boundaries. The genes containing WRE or WRE+ or none are listed in Table S6. The code for the identification of C-clamp targets is available at https://zenodo.org/records/24375870.

We intersected the genes containing the WRE+ motif with all the genes expressed in iPSC at DE 4h (Table S4). We determined the differentially expressed genes between Ctrl and *CTNNB1* Udel cells (log2FC >1.5) and divided them equally into genes strongly or weakly regulated in Udel cells.

### Gene ontology analysis

Gene ontology analysis was performed using DAVID^108^. Bonferroni-adjusted *P* values are reported.

### Statistical analysis

Statistical parameters are reported in the figures and figure legends, including the definitions and exact values of *N* and experimental measures. Transcriptome-wide feature comparisons were performed using a two-sided Mann-Whitney test, a Wilcoxon test or a contingency test in SPSS. For experimental data, statistical analyses were performed using Welch’s *t*-test to compare differences between two groups, unless otherwise specified in the figure legends. Data are presented as mean ± SD or mean ± SEM. Statistical significance is indicated as follows: ns: not significant, *, *P* < 0.05; **, *P <* 0.01, ***, *P* < 0.001, and ****, *P <* 0.0001. Boxplots depict median, 25^th^ and 75^th^ percentile (boxes) and 5% and 95% confidence intervals (error bars).

---

**Figure S1.**
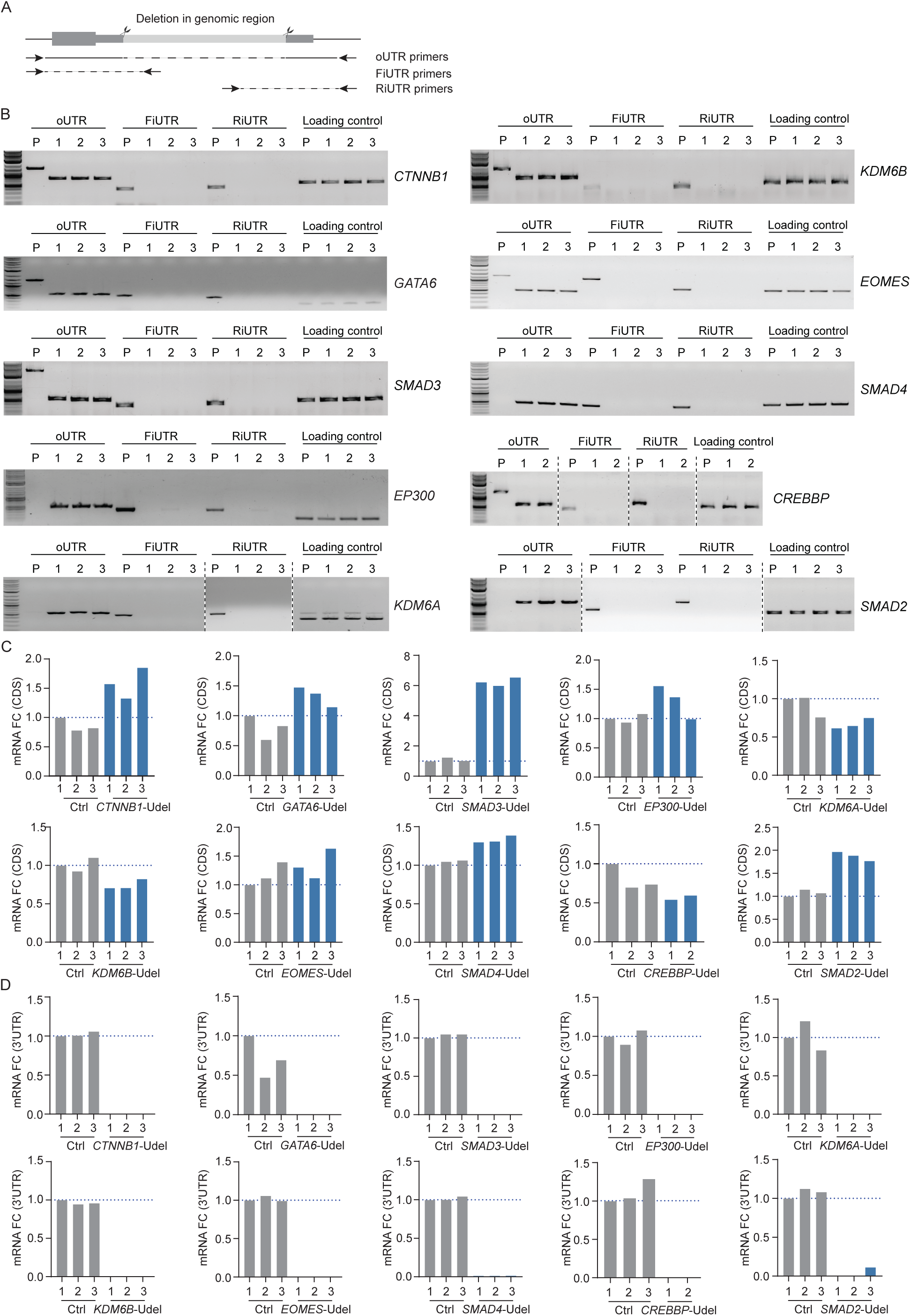
Validation of 3′UTR deletions at the DNA and mRNA level, related to. **Figure 1. A.** PCR primer design used to examine genomic deletions that disrupt mRNA 3′UTRs. oUTR primers flank the 3′UTR region, whereas FiUTR and RiUTR primer pairs span the junction between the genome and the deleted region. **B.** Representative gel images of validation of 3′UTR deletions at the DNA level using PCR. P, parental cells, numbers indicate clonal lines of the respective 3′UTR deletions. Amplification of an unedited region within the *MYC* locus serves as loading control. **C.** mRNA transcript expression, measured in the coding sequence (CDS) in Ctrl and Udel clones. *GAPDH* was used for normalization. **D.** As in (C), mRNA transcript expression was measured with primers located in the indicated 3′UTRs.

**Figure S2.**
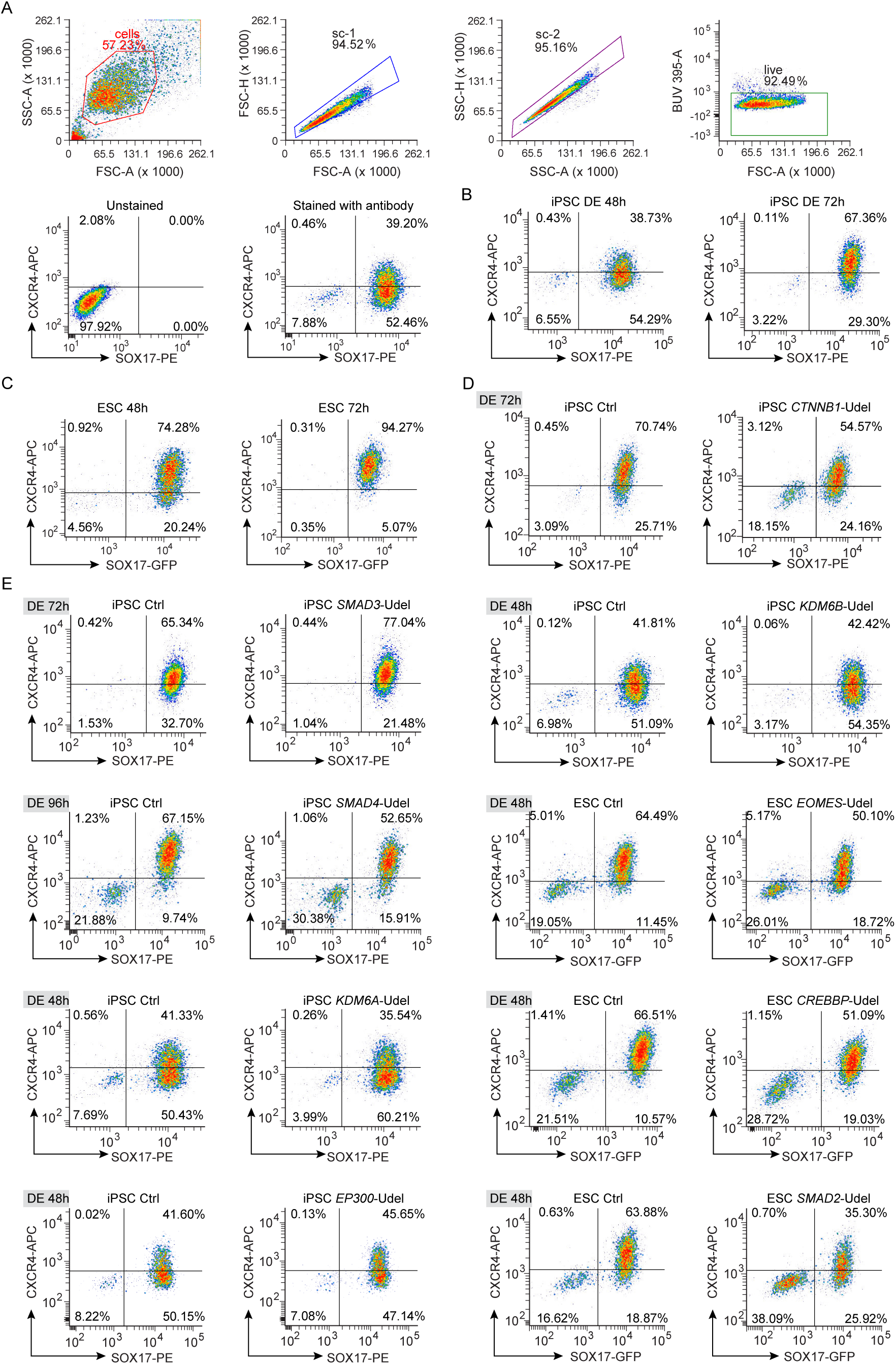
Endogenous 3′UTR deletions alter DE differentiation efficiency in pluripotent stem cells, related to. **Figure 1. A.** Representative flow plots showing the gating strategy to identify DE cells. Cells were first gated with FSC-A and SSC-A, followed by gating using FSC-A/FSC-H and SSC-A/SSC-H, and then gated for the live-cell population. The live-cell gate was used for SOX17 and CXCR4 analysis. Representative plots of SOX17 and CXCR4-stained cells and unstained control cells at DE 48h are shown. **B.** Representative flow plots showing the DE differentiation efficiency, assessed by SOX17+CXCR4+ double-positive cells at 48h or 72h post-DE induction in iPSCs or embryonic stem cells (ESCs; HUES8), respectively. **C.** As in (B), but shown for HUES8 ESCs. **D.** As in (B), but shown for Ctrl and CTNNB1 Udel clone at 72h post-DE induction. **E.** As in (D), but shown for Ctrl and indicated Udel clones at the indicated timepoints. Shown is the earliest timepoint at which a significant change in DE differentiation efficiency was observed in Udel cells. For *SMAD3* and *SMAD4* Udel clones, a significant difference in SOX17+CXCR4+ double-positive cells was observed at DE 72h and DE 96h, respectively.

**Figure S3.**
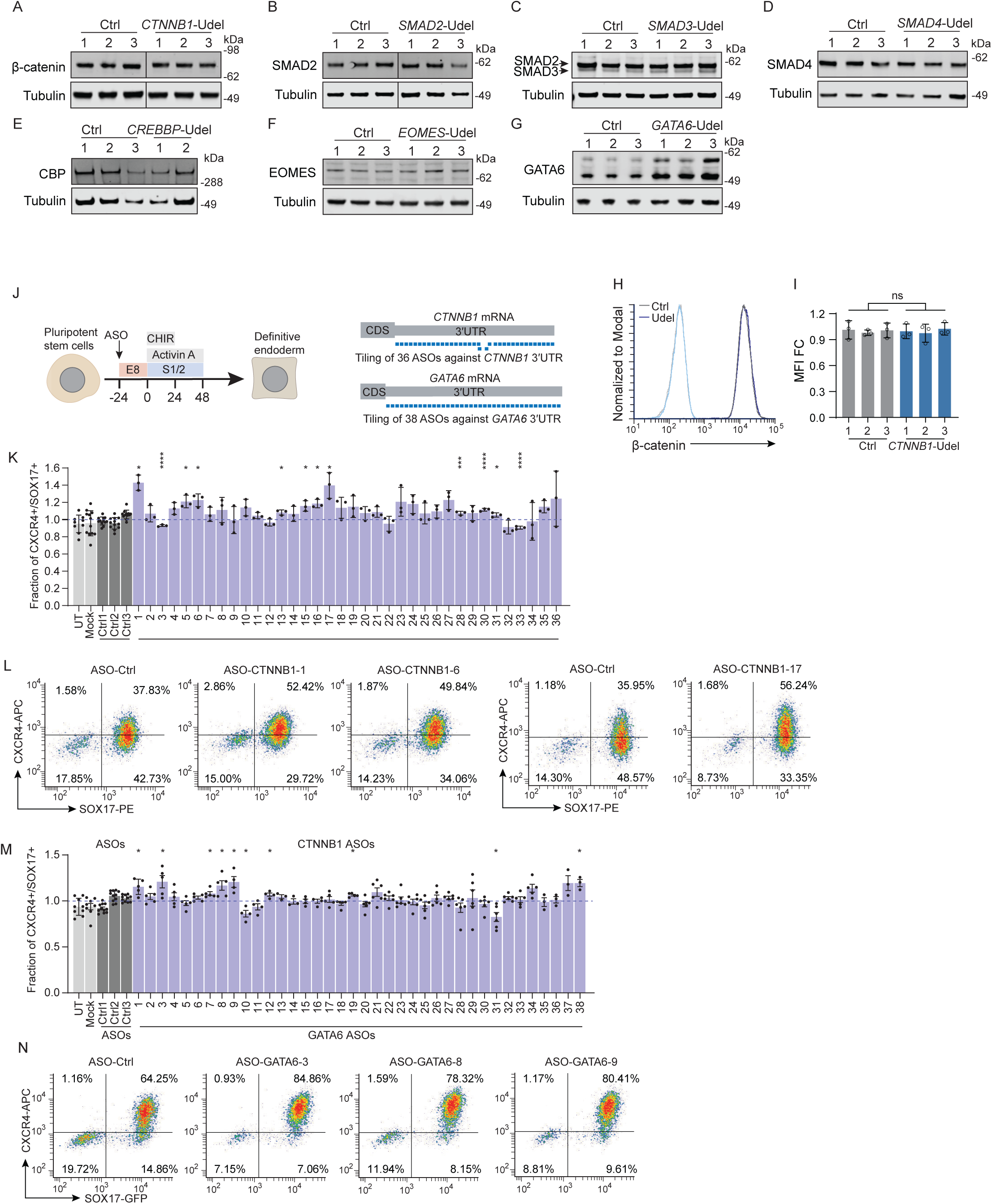
Lack of protein abundance change in most 3′UTR deletion cells and blocking of 3′UTRs through ASOs, related to. **Figure 1. A.** Immunoblots showing β-catenin protein abundance in Ctrl and *CTNNB1* Udel cells at the stem cell stage. Shown are 3 clonal lines. Tubulin serves as loading control. **B.** As in (A) but shown for SMAD2 in Ctrl and *SMAD2* Udel cells. **C.** As in (A) but shown for SMAD3 in Ctrl and *SMAD3* Udel cells. An antibody against SMAD2/3 was used. The bottom band refers to SMAD3. **D.** As in (A) but shown for SMAD4 in Ctrl and *SMAD4* Udel cells. **E.** As in (A) but shown for CBP in Ctrl and *CREBBP* Udel cells. *N* = 2 clonal lines. **F.** As in (A) but shown for EOMES in Ctrl and *EOMES* Udel cells at DE 24h. **G.** As in (A) but shown for GATA6 in Ctrl and *GATA6* Udel cells at DE 24h. **H.** Representative flow plots showing β-catenin protein abundance in Ctrl and Udel iPSCs. **I.** Quantification of the experiment from (H). β-catenin mean fluorescence intensity (MFI) is shown as mean ± SD from 3 independent experiments from 3 independent clonal lines. Welch’s *t*-test; ns, not significant. **J.** Schematic of ASO transfection followed by DE differentiation. Schematic of tiling of 36 and 38 ASOs across the entire *CTNNB1* and *GATA6* 3′UTRs, respectively. **K.** Quantification of SOX17+CXCR4+ double-positive cells by flow cytometry at DE 48h. Each experiment included WT iPSCs that were untreated (UT), transfected with RNAiMAX only (mock), or transfected with control ASOs (Ctrl 1–3). Each CTNNB1 ASO was transfected individually. In each ASO experiment, values obtained by the CTNNB1 ASOs were normalized to the mean of the three Ctrl ASOs. A total of 12 independent experiments was performed. Statistical analysis was conducted using Welch’s *t* test, comparing each ASO with all the values obtained from the three Ctrl ASOs. Data are shown as mean ± SD. Welch’s *t*-test; *, *P*< 0.05; ***, *P*< 0.001; ****, *P*< 0.0001. **L.** Representative flow plots showing the DE differentiation efficiency at 48h post-induction in WT cells transfected with individual ASOs against the *CTNNB1* 3′UTR. **M.** As in (K), but HUES8 cells and ASOs against the *GATA6* 3′UTR were used. **N.** As in (L), but individual GATA6 ASOs are shown.

**Figure S4.**
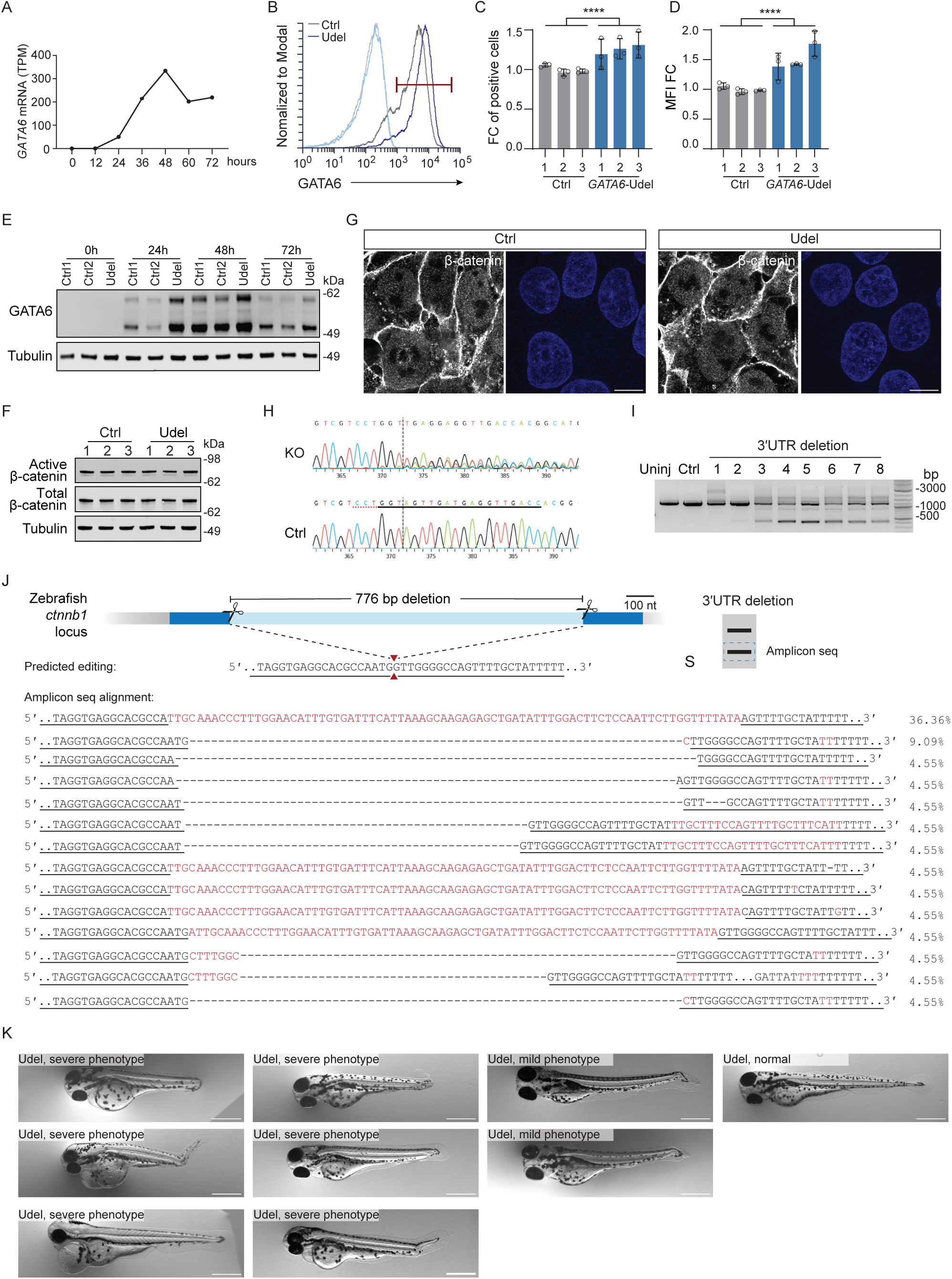
Deletion of the *ctnnb1* 3′UTR in zebrafish, related to. **Figure 1 and 2. A.** *GATA6* mRNA expression shown as TPM from published single-cell RNA-seq data^34^. **B.** Representative flow cytometry plots of GATA6 in control (dark gray) and *GATA6* Udel (dark blue) cells at 24h of DE differentiation. Unstained samples are shown in lighter colors. **C.** Quantification of the experiment from (B). Shown is mean ± SD from 3 independent experiments from 3 independent clonal lines. Welch’s *t*-test; ****, *P* < 0.0001. **D.** As in (C) but MFI of GATA6-positive cells is shown. **E.** Immunoblot showing GATA6 protein abundance at the indicated timepoints. Tubulin serves as loading control. **F.** Immunoblot showing active and total β-catenin in Ctrl and Udel cells at DE 24h. *N* = 3 clonal lines were examined. Tubulin serves as loading control. **G.** Representative immunofluorescence images showing β-catenin distribution at DE 4h. DAPI was used to stain the nucleus. Scale bar, 10 μm. Quantification is shown in Fig. 2G. **H.** Sanger sequencing of edited alleles of zebrafish *ctnnb1* KO embryos. The guide RNA is underlined, and the cut site is indicated by the vertical dashed line. **I.** Representative gel image of genotyping PCR to validate zebrafish *ctnnb1* 3′UTR deletions using deletion-flanking primers. **J.** Schematic of the zebrafish *ctnnb1* 3′UTR deletion region. The predicted post-editing sequence is shown. Red arrowheads indicate the junction site after scarless repair. In mosaic embryos, edited PCR products were excised and analyzed by amplicon sequencing. Sequences were aligned and composition is shown. Sequences matching the prediction are underlined; insertions are highlighted in red. **K.** As in Fig. 2K, but additional zebrafish images are shown.

**Figure S5.**
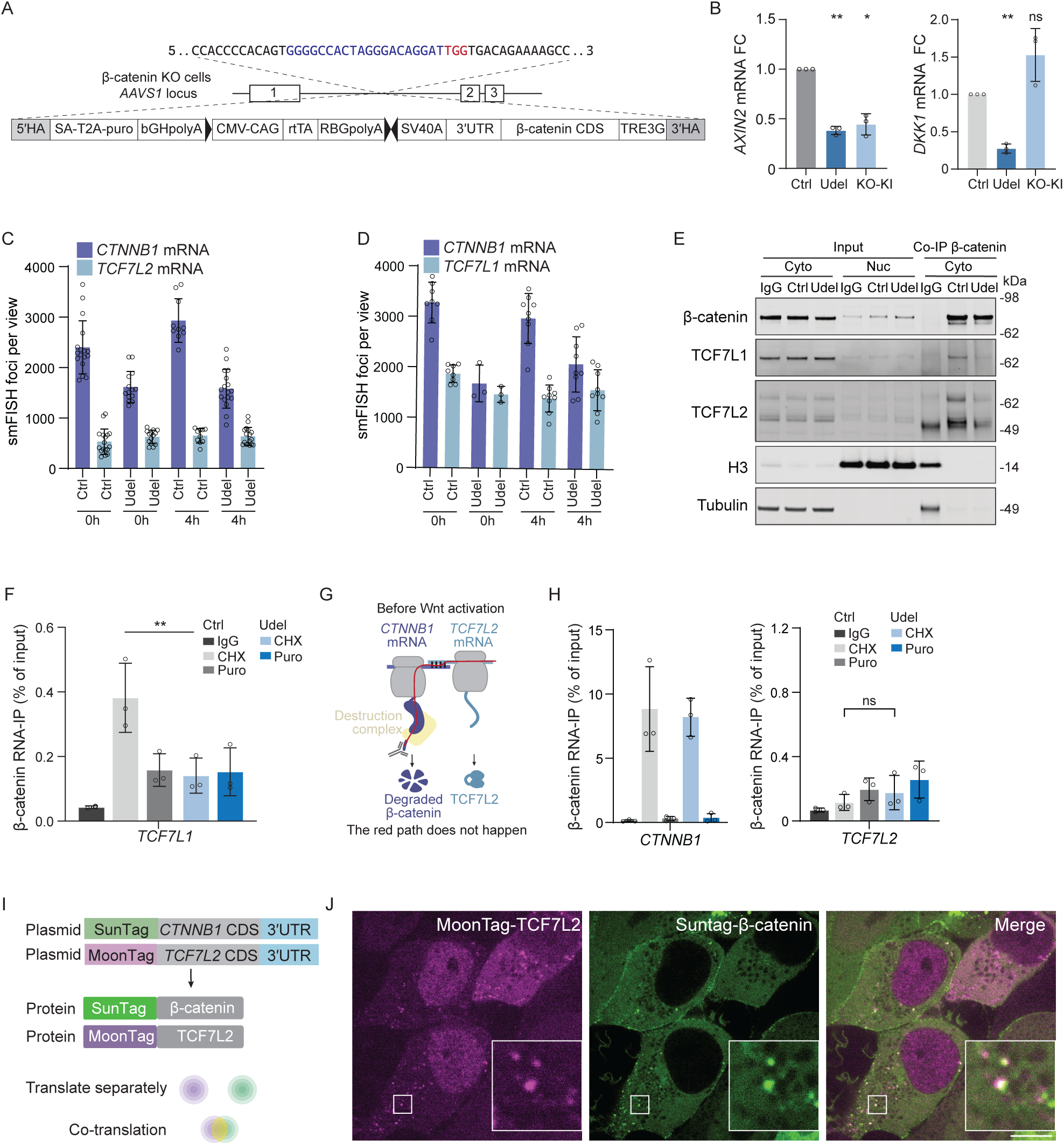
Wnt signaling triggers mRNA heterodimerization-driven cofolding of β-catenin and TCF7L2, related to. **Figure 3 and 4. A.** Schematic of the KI construct used for rescue experiment performed in *CTNNB1* KO iPSCs. **B.** As in Fig. 3C, but *AXIN2* and *DKK1* mRNA FC is shown. Welch’s *t*-test. **, *P* < 0.0024; *, *P* = 0.012. **C.** Absolute numbers of *CTNNB1* and *TCF7L2* smRNA-FISH foci per field of view in Ctrl and Udel cells, at stem cell stage (0h) and at DE 4h. Each dot represents one field of view including 9-15 cells. Shown is mean ± SD from 2 independent experiments. **D.** As in (C), but *CTNNB1* and *TCF7L1* are shown. **E.** Co-IP of endogenous β-catenin with endogenous interactors at DE 4h in the cytoplasmic fraction. 2% of the cytoplasmic fraction was used as input. Nuclear and cytoplasmic fractions were loaded at 1:1 ratio. Tubulin serves as loading control for the cytoplasmic fraction, whereas H3 serves as loading control for the nuclear fraction. Reduced co-IP of TCF7L1 and TCF7L2 in Udel compared with Ctrl cells. **F.** As in Fig. 4M, but β-catenin RNA-IP followed by *TCF7L1* qRT-PCR is shown. Welch’s *t*-test. *, *P* = 0.041 **G.** Schematic of *CTNNB1* and *TCF7L2* mRNA heterodimerization and co-translational protein complex formation before Wnt signaling activation. The experiment shown in (H) rules out the red path. **H.** β-catenin RNA-IP at the stem cell stage followed by *CTNNB1* and *TCF7L2* qRT-PCR. Shown is mean ± SD from 3 independent experiments. Welch’s *t*-test; ns, not significant. **I.** Schematic of SunTag-β-catenin (green) and MoonTag-TCF7L2 (magenta) translation reporters. Separate translation yields distinct SunTag and MoonTag foci, whereas co-translation yields co-localization signal. **J.** Representative frame from live-cell imaging of co-translating SunTag-β-catenin and Moontag-TCF7L2 after treatment with 5 µM CHIR99021 for 2h in HEK293T cells. The foci represent individual β-catenin (green) and TCF7L2 (magenta) protein molecules, whereas yellow/white foci represent co-localized β-catenin and TCF7L2. Scale bar, 10 μm.

**Figure S6.**
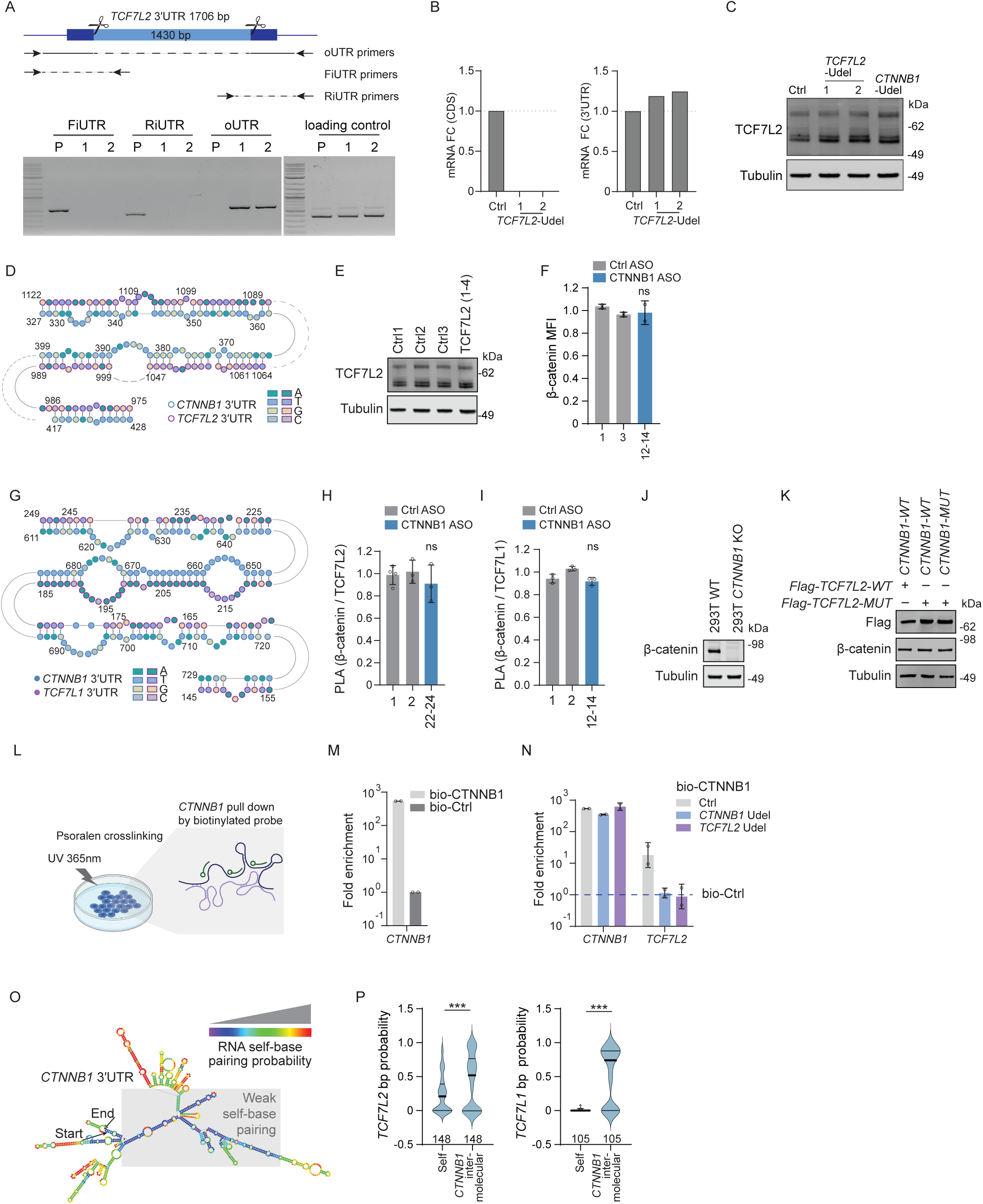
The intermolecular 3′UTR-3′UTR interaction between *CTNNB1* and *TCF7L2* is direct, related to. **Figure 5. A.** Schematic of PCR primer design to examine the *TCF7L2* 3′UTR deletion (top). Representative gel images of PCR results in parental (P) cells and *TCF7L2* Udel clones (*N* = 2 clonal lines) are shown. Amplification of the unedited region within the *MYC* locus serves as loading control. **B.** As in Fig. S1C and S1D but shown is mRNA transcript abundance of *TCF7L2* mRNA. Shown are 2 clonal lines. **C.** Immunoblot of TCF7L2 protein in Ctrl and *TCF7L2* Udel iPSCs at the iPSC stage. Tubulin serves as loading control. **D.** The intermolecular *CTNNB1-TCF7L2* 3′UTR-3′UTR interaction according to RNA bifold is shown. The numbers indicate the nt of the respective 3′UTRs. **E.** Immunoblot analysis of TCF7L2 protein in Ctrl and TCF7L2 ASO treated cells at DE 4h. Tubulin serves as loading control. **F.** β-catenin protein quantified by flow cytometry at DE 4h after addition of Ctrl ASOs or CTNNB1 ASOs 12-14. MFI is shown as mean ± SD from 2 independent experiments. Welch’s *t*-test; ns, not significant. **G.** As in (D), but shown for the *CTNNB1* and *TCF7L1* 3′UTRs. **H.** As in Fig. 5F, but CTNNB1 ASOs 21-24 are shown. Welch’s *t*-test. ns, not significant. **I.** As in Fig. 5H, but CTNNB1 ASOs 12-14 are shown. Welch’s *t*-test. ns, not significant. **J.** Immunoblot of β-catenin in WT and *CTNNB1* KO HEK293T cells. Tubulin serves as loading control. **K.** Immunoblot of β-catenin and Flag-tagged TCF7L2. *CTNNB1* KO HEK293T cells stably expressing the *CTNNB1* CDS and WT or mutant (MUT) 3′UTR were transiently transfected with Flag-tagged TCF7L2 CDS containing WT or MUT 3′UTR. Protein expression was analyzed in cells treated with 3 µM CHIR99021 for 6h. Tubulin serves as loading control. **L.** Schematic of psoralen-based chemical crosslinking, followed by mRNA pull-down to detect direct mRNA-mRNA interactions in iPSCs. Biotinylated probes against *CTNNB1* mRNA are shown in green. **M.** Fold enrichment of *CTNNB1* mRNA pulled down using biotinylated CTNNB1 probes relative to biotinylated control probes against *TMSB4X* mRNA. **N.** Fold enrichment of pulled down *CTNNB1* and *TCF7L2* mRNAs using biotinylated CTNNB1 probes relative to control probes in Ctrl, *CTNNB1* Udel, and *TCF7L2* Udel clones. Shown is mean ± SD from 2 independent experiments. **O.** RNAfold output of MFE structure of *CTNNB1* 3′UTR, color-coded by RNA self-base pairing probability. **P.** Left panel, base pairing probability of the *TCF7L2* dimerization site is shown for *TCF7L2* self-interaction and *CTNNB1-TCF7L2* intermolecular interaction. Wilcoxon test, *P* = 3.7 x 10^−7^. Right panel, base pairing probability of the *TCF7L1* dimerization site is shown for *TCF7L1* self-interaction and *CTNNB1-TCF7L1* intermolecular interaction. Wilcoxon test, *P* = 8.1 x 10^−13^.

**Figure S7.**
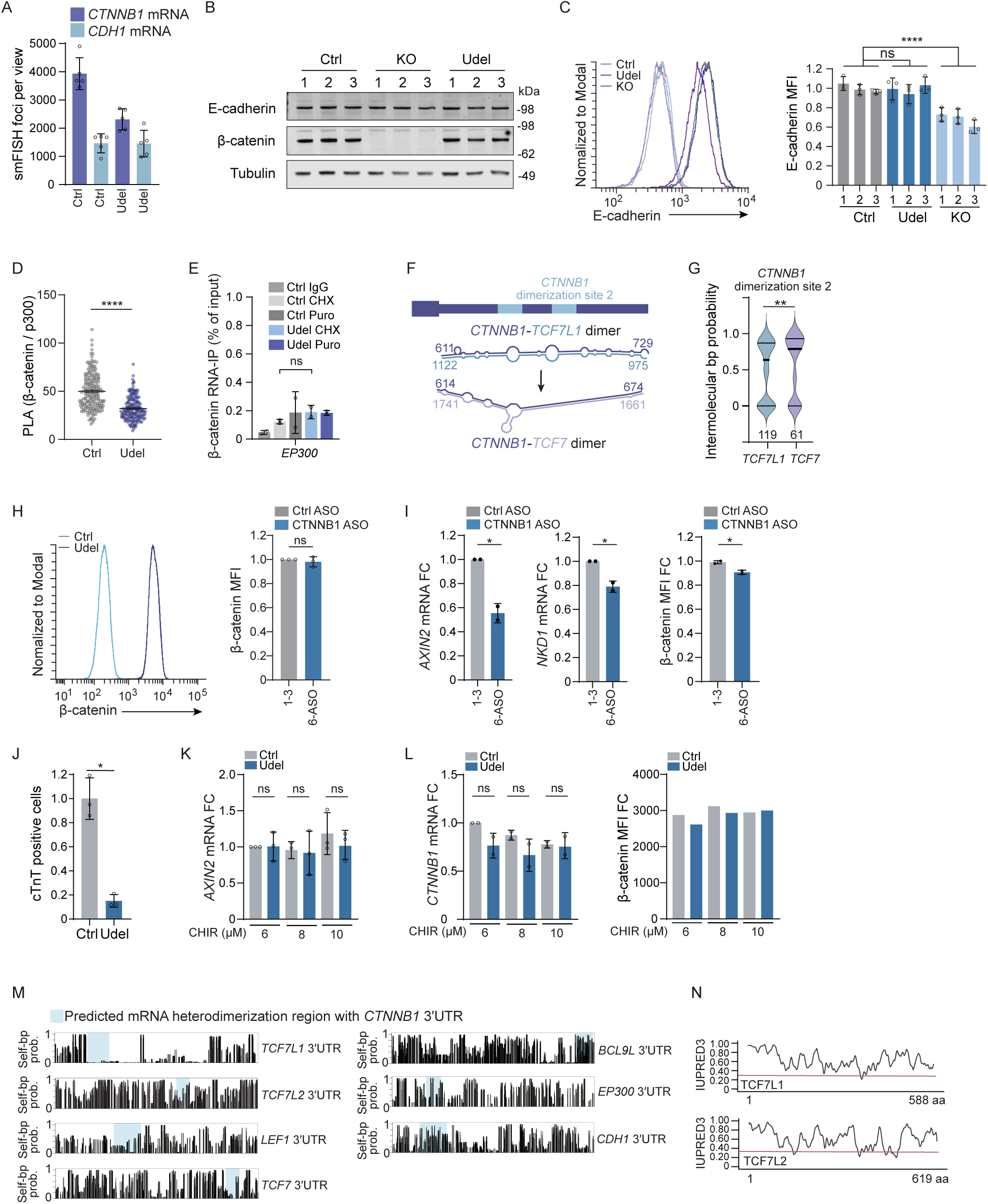
*CTNNB1* mRNA dimerizes with Wnt transcription factor mRNAs but not with *CDH1* mRNA, related to. **Figure 6 and 7. A.** Absolute numbers of *CTNNB1* and *CDH1* smRNA-FISH foci per field of view in Ctrl and Udel cells, at stem cell stage. Each dot represents one field of view including 9-15 cells. Shown is mean ± SD from of one experiment out of 2 independent experiments. **B.** Immunoblot of total E-cadherin and β-catenin protein in iPSCs. *N* = 3 clonal lines each were used, and Tubulin serves as loading control. **C.** Left panel, representative flow plots showing surface E-cadherin in control (dark gray), *CTNNB1* KO (dark purple) and *CTNNB1* Udel (dark blue) cells at the iPSC stage. The unstained samples are shown in lighter colors. Right panel, quantification of surface E-cadherin by MFI, shown as mean ± SD from 3 independent experiments of 3 clonal lines, each. Welch’s *t*-test. ****, *P* = 2.5 x 10^−7^. **D.** Quantification of β-catenin/p300 PLA foci at DE 4h. Cells analyzed, Ctrl, *N* = 182; Udel, *N* = 174. Shown is mean ± SEM from 2 independent experiments. Welch’s *t*-test. ****, *P* = 1.65 x 10^−23^. **E.** β-catenin RNA-IP from Fig. 4M, followed by qRT-PCR for *EP300* mRNA. Shown is mean ± SD from 2 independent experiments. Welch’s *t*-test; ns, not significant. **F.** RNA bifold prediction of mRNA heterodimerization between the *CTNNB1* and *TCF7* 3′UTRs, which occurs in the *CTNNB1* dimerization site 2. This region is also the binding site of the *TCF7L1* 3′UTR. Shown as in Fig. 6K. **G.** Base pairing strength in the *CTNNB1* dimerization site 2 upon intermolecular interaction with the *TCF7L1* or *TCF7* 3′UTRs. Wilcoxon test, *P* = 0.0002. **H.** β-catenin protein quantified by flow cytometry at DE 4h. A representative flow plot is shown, with unstained samples in lighter colors. Right panel, β-catenin MFI as mean ± SD from 3 independent experiments. Welch’s *t*-test; ns, not significant. **I.** As in Fig. 6O, but experiments were performed in HCT116 cells. *AXIN2* and *NKD1* mRNA levels were examined, and β-catenin protein abundance levels were quantified by flow cytometry. Shown is mean ± SD from 2 independent experiments. *T*-test, *, *P* < 0.05. **J.** Cardiomyocyte differentiation efficiency was measured by the fraction of cardiac troponin T (cTnT)-positive cells at d12. Shown is mean ± SD from 3 independent experiments. Welch’s *t*-test. **, *P* = 0.009. **K.** As in Fig. 7E, but *AXIN2* mRNA FC is shown. Welch’s *t*-test; ns, not significant. **L.** As in Fig. 7E, but *CTNNB1* mRNA FC is shown. Welch’s *t*-test; ns, not significant. Right panel, β-catenin protein was quantified by flow cytometry. **M.** RNA self-base pairing probablity for each 3′UTR nt is shown for the indicated 3′UTRs. The predicted binding sites of the *CTNNB1* 3′UTR are highlighted in light blue. **N.** IUPRED3 profiles of TCF7L1and TCF7L2 showing that they predicted to be predominantly disordered.

